# Conformational catalysis of cataract-associated aggregation in a human eye lens crystallin occurs via interface stealing

**DOI:** 10.1101/601708

**Authors:** Eugene Serebryany, William M. Jacobs, Rostam M. Razban, Eugene I. Shakhnovich

## Abstract

Human γD-crystallin (HγD) is an abundant and highly stable two-domain protein in the core region of the eye lens. Destabilizing mutations and post-translational modifications in this protein are linked to onset of aggregation that causes cataract disease (lens turbidity). Wild-type HγD greatly accelerates aggregation of the cataract-related W42Q variant, without itself aggregating. The mechanism of this “inverse prion” catalysis of aggregation remained unknown. Here we provide evidence that an early unfolding intermediate with an opened domain interface enables transient dimerization of the C-terminal domains of wild-type and mutant, or mutant and mutant, HγD molecules, which deprives the mutant’s N-terminal domain of intramolecular stabilization by the native domain interface and thus accelerates its misfolding to a distinct, aggregation-prone intermediate. A detailed kinetic model predicts universal power-law scaling relationships for lag time and rate of the resulting aggregation, which are in excellent agreement with the data. The mechanism reported here, which we term interface stealing, can be generalized to explain how common domain-domain interactions can have surprising consequences, such as conformational catalysis of unfolding, in multidomain proteins.

**Significance:** Most known proteins in nature consist of multiple domains. Interactions between domains may lead to unexpected folding and misfolding phenomena. This study of human γD-crystallin, a two-domain protein in the eye lens, revealed one such surprise: conformational catalysis of misfolding via intermolecular domain interface “stealing.” An intermolecular interface between the more stable domains outcompetes the native intramolecular domain interface. Loss of the native interface in turn promotes misfolding and subsequent aggregation, especially in cataract-related γD-crystallin variants. This phenomenon is likely a contributing factor in the development of cataract disease, the leading worldwide cause of blindness. However, interface stealing likely occurs in many proteins composed of two or more interacting domains.

## Introduction

Beyond the native and fully unfolded states, most proteins can adopt partially unfolded or misfolded intermediates. Intermediates are especially important for multidomain proteins, which comprise most eukaryotic proteins yet only a small minority of biophysical studies to-date (1–3). These intermediates are often rare or transient and thus challenging to investigate, yet they have large effects on the protein’s stability, evolution, and function. Misfolded intermediates can dominate folding pathways (4–7), drive evolution of the coding sequence (8–10), and cause proteins to assemble into aggregates linked to common diseases (11–17). Post-translational modifications often affect the relative stabilities of intermediates within the conformational ensemble (18–24). Where a protein is present at a high concentration, the question arises: Can distinct intermediate states interact or affect one another in a functionally significant way? Here we present experimental evidence that domain interface opening in one molecule of a stable two-domain protein can catalyze domain misfolding in another.

Vertebrate lens crystallins are extremely long-lived; in the core region of the eye lens they are synthesized *in utero* and never replaced thereafter (25–27). Human γD-crystallin is extremely thermodynamically and kinetically stable, likely an evolutionary adaptation to resist aggregation (17, 25, 28–30). However, crystallins accumulate damage throughout life, even as the cytoplasm of aged lens cells becomes increasingly oxidizing (31–38). Eventually, aggregation-prone partially unfolded intermediates arise, notably those locked by non-native disulfides (35, 39–45). Cataract is lens opacity that results when these misfolded crystallins assemble into light-scattering aggregates, making the lens turbid. Cataract is a disease of aging, affecting 17% of all people aged 40 and over (46, 47), though many congenital or early-onset mutations are also known (48–50). Due to the high cost of surgery and lack of a therapeutic treatment, cataract remains the leading cause of blindness in the world (47, 51).

HγD consists of two homologous double-Greek key domains joined by a non-covalent domain interface (52, 53). This interface contributes significantly to the stability of the N-terminal domain (53–55). Destabilization and misfolding of the N-terminal domain under oxidizing conditions leads to an aggregation-prone intermediate state kinetically trapped by a non-native internal disulfide bond between Cys32 and Cys41 (43, 56). Importantly, formation of the same non-native disulfide is supported by tissue proteomics of aged cataractous lenses even in the absence of any crystallin mutations (35). This observation suggests that multiple modes of damage – destabilizing core mutations, post-translational modifications, or possibly even thermal fluctuations – converge on the same misfolded aggregation precursor. Surprisingly, the highly stable wild-type HγD (WT) promotes aggregation of its destabilized W42Q variant without itself aggregating – an interaction that may produce significant lens turbidity from even a minor fraction of damaged molecules (57).

We have previously reported the crucial interplay between misfolding and disulfide exchange in the aggregation of cataract-associated HγD variants (44). Specifically, we showed that HγD molecules can exchange disulfide bonds dynamically in solution, which leads to the “hot potato” model: if a molecule with a destabilized N-terminal domain accepts a disulfide bond from a more stable molecule, misfolding and aggregation of the destabilized variant will ensue (44).

We now demonstrate that the wild-type protein also promotes aggregation of multiple cataract-related variants of HγD by a novel conformational mechanism, active even when the redox function is abolished. This mechanism is associated with opening of the domain interface in the WT protein. Population of this interface-opened intermediate enables a transient “interface stealing” interaction between WT and mutant molecules such that the N-terminal domain of the aggregation-prone variant loses its interface with the C-terminal domain and hence the kinetic and thermodynamic stabilization derived from that interface. Thus, the interaction facilitates unfolding of the mutant’s N-terminal domain leading to its conformational transformation and subsequent aggregation. We refer to this process as conformational catalysis.

## Results

### Conformational catalysis of W42Q, L5S, and V75D aggregation

The W42Q variant of HγD mimics UV-induced damage to a Trp residue and shows significant propensity to aggregate at physiological temperature and pH, similar to the congenital-cataract mutant W42R (43, 45, 58). This aggregation process requires formation of a non-native internal disulfide bond (Cys32-Cys41), which traps the protein in a non-native conformation characterized by extrusion of the N-terminal β-hairpin (43). Because of this requirement of oxidation combined with misfolding, aggregation does not occur in the absence of oxidizing agent (43, 44).

When the oxidizing agent glutathione disulfide (GSSG) was present in the buffer, aggregation of W42Q proceeded readily at 37 °C and pH 7 in a concentration-dependent manner (Fig. 1A). We have previously shown that GSSG can substitute for a disulfide in wild-type HγD (Cys108-Cys110) as the oxidizing agent for W42Q aggregation (44). We investigated whether presence of WT produced any further effect on W42Q aggregation when excess GSSG was also present in the buffer. Addition of WT to W42Q+GSSG led to dramatic, [WT]-dependent acceleration of W42Q aggregation despite the excess GSSG (Fig. 1B). Adding more GSSG did not significantly affect aggregation (Fig. 1C). Removing any Cys residue from the WT protein by site-directed mutagenesis did not abolish WT-induced acceleration of the aggregation rate, and often actually enhanced it (Fig. 1D). Furthermore, it did not matter whether the WT protein was oxidized or reduced – as long as a source of disulfides was present in excess, aggregation in the WT/W42Q mixture was rapid (Fig. S1). These findings indicated that there exists a distinct mechanism by which WT promotes aggregation of W42Q independent of its oxidoreductase activity.

To confirm that the increase in solution turbidity was due to increased W42Q aggregation and not to co-aggregation of WT and W42Q, we determined the composition of the aggregated fraction. To that end we used a previously established PEGylation assay (44) to determine the total number of Cys residues per protein molecule in soluble vs. aggregated fractions of W42Q mixed with one of two multi-Cys mutants in the WT background. The variants used were C18T/C41A/C78A (“CCC”) and C18T/C78A/C108S/C110S (“CCCC”). Like WT, both variants promoted W42Q aggregation (Fig. 1D). Importantly, both CCC and CCCC variants remained entirely within the soluble supernatant fraction at the end of the aggregation reaction, while the majority of W42Q transferred to the aggregated fraction (Fig. 1E). All samples were reduced prior to PEGylation, hence W42Q reacted with 6 molecules of PEG-maleimide per protein molecule; a minor band at 4 PEG-maleimide per protein molecule was evidence of incomplete reduction, likely due to burial of the 32-SS-41 disulfide in condensed aggregates. As a complementary experimental approach, we examined the composition of soluble and aggregated fractions in W42Q/WT mixtures by MALDI-TOF mass spectrometry, which likewise showed only W42Q in the aggregated fraction and WT in the supernatant (Fig. S2). We conclude that the increase in solution turbidity in W42Q/WT or W42Q/CCC or W42Q/CCCC mixtures was not due to co-aggregation, but rather to increased aggregation of the W42Q variant. This is consistent with initial work on this system (57).

**Fig. 1:**
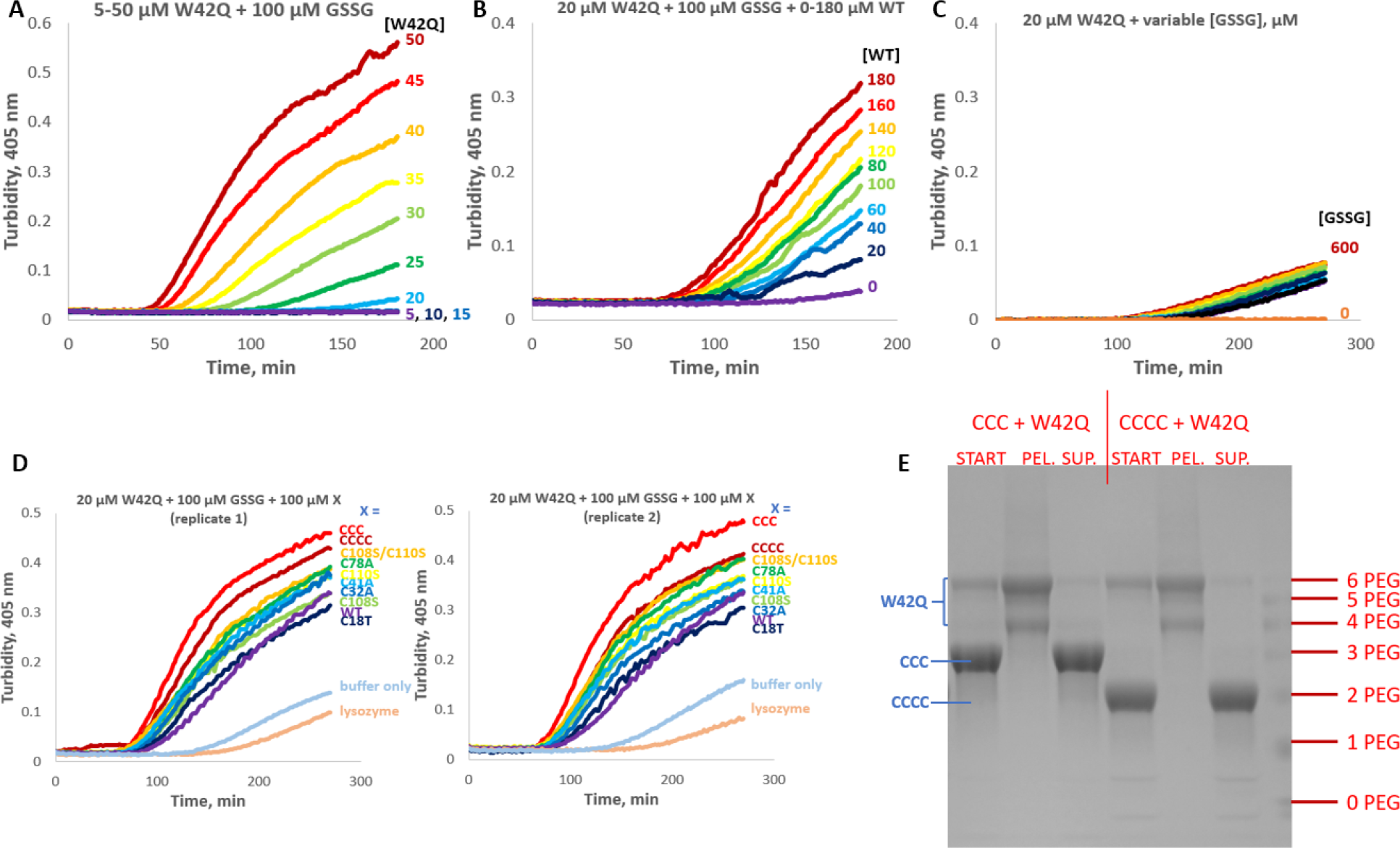
WT HγD promotes aggregation of the W42Q variant in a catalytic, redox-independent manner. (A) W42Q HγD aggregated in a concentration-dependent manner in the presence of excess glutathione disulfide (GSSG). (B) WT promoted aggregation of W42Q in a [WT]-dependent manner. (C) Further excess of GSSG did not significantly affect W42Q aggregation. (D) Mutagenesis of any given Cys residue, or the combination of C18T/C41A/C78A (“CCC”) or C18T/C78A/C108S/C110S (“CCCC”) in the WT background did not abolish its ability to promote W42Q aggregation at saturating GSSG, and indeed enhanced it. A control protein (lysozyme) did not promote W42Q aggregation. Two replicates are shown as illustration of reproducibility. (E) Cys-counting analysis of the starting (“START”), aggregated (“PEL”), and supernatant (“SUP”) fractions of W42Q/CCC and W42Q/CCCC mixtures. The W42Q variant contains 6 Cys residues and thus reacted with a maximum of 6 PEG-maleimide equivalents (4 if the internal disulfide required for aggregation has not been reduced). The CCC and CCCC variants contain only 3 and 2 Cys residues, respectively. Neither CCC nor CCCC was detectable in the aggregated fraction, which was composed entirely of W42Q, indicating that the aggregation-promoting activity was catalytic.

To assess the generality of the findings beyond W42Q, we tested two other variants of HγD: V75D and L5S. Even though the three mutations are in distal parts of the N-terminal domain’s sequence, all three are associated with cataract (43, 48, 59–61), and all three replace core hydrophobic residues with hydrophilic ones, thus destabilizing the N-terminal domain (see Table 1). Each variant showed redox-dependent aggregation comparable to that of W42Q, although at somewhat higher protein concentrations (Fig. S3). This difference will be explained in the structural and kinetic model sections below. The convergence among W42Q, L5S, and V75D established that conformational catalysis is not due to a unique feature of the W42Q variant. The common feature among all three variants is simply the reduced stability of the N-terminal domain core.

To directly measure whether presence of a catalytic variant accelerates unfolding of an aggregation-prone mutant’s N-terminal domain, we used time-resolved PEGylation of a mixture of W42Q and CCCC using a maleimide-PEG reagent. Since the N-terminal domain of W42Q contains four Cys residues, three of which are buried in the domain core and the fourth in the domain interface, Cys accessibility is a good probe for foldedness of the domain. Samples from W42Q and a W42Q/CCCC mixture incubated in the presence of PEG-maleimide at 37 °C in native buffer were taken at fixed intervals, the PEGylation reagent quenched with free cysteine, and the proteins resolved by SDS-PAGE to quantify the distribution of singly and multiply PEGylated molecules (Fig. 2 and Fig. S4).

The CCCC sample contains only two Cys residues, one buried in the N-terminal core and the other in the domain interface. In control samples of CCCC alone only one Cys residue was found to be reactive during an hour-long incubation (Fig. 2, *gray*). We assign this to the residue buried in the domain interface, Cys41; the kinetics of its reactivity is related to the kinetics of interface opening in this variant.

By contrast, the W42Q protein reacted extensively with PEG-maleimide during the course of the incubation, and both initial and full PEGylation was achieved more rapidly in the W42Q/CCCC mixture than in W42Q alone (Fig. 2). The first reactive site (+1 PEG) is assigned to Cys110, which is the only natively exposed and the most reactive thiol (37, 44, 52). The second reactive site in W42Q (+2 PEG) may be Cys41, since its solvent exposure requires only opening of the domain interface, which occurs relatively easily (43, 55, 62). However, no definitive assignment has been made. Kinetics of reactivity at the second site, as well as for full unfolding (+6 PEG) were significantly faster in the presence vs. absence of the conformational catalyst. Flux through the intervening numbers of exposed thiols was comparable, however, suggesting that following exposure of the first (or possibly second) natively buried Cys residue unfolding becomes cooperative. We conclude that the aggregation-catalyzing interaction between W42Q and CCCC has a structural basis: it increases the unfolding rate of the W42Q N-terminal domain. Consistent with this, prior work has revealed that WT downshifts the aggregation-permissive temperature for the mutant (57).

**Fig. 2:**
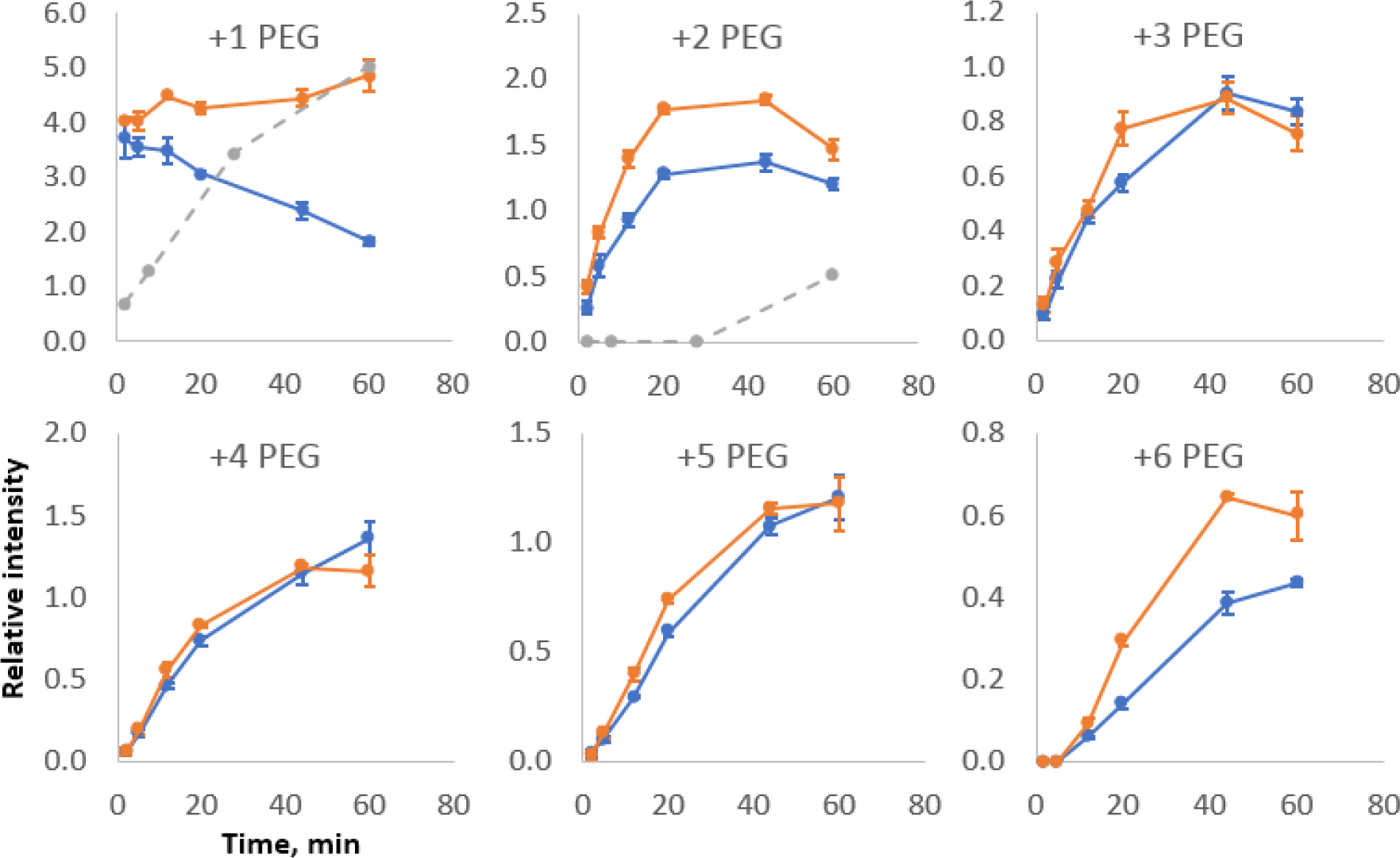
Presence of a catalytic variant accelerates W42Q unfolding. W42Q unfolded more rapidly when CCCC was present (*orange*) than without CCCC (*blue*), as reported by PEGylation, a measure of solvent accessibility of buried Cys residues. Error bars show S.E.M. of 3-4 replicate measurements with independent quantitation by SDS-PAGE. Control experiments with CCCC alone show the reactivity of its only two Cys residues (*gray*). PEGylation gels are shown in Fig. S3.

### Structural model of conformational catalysis of HγD aggregation

The finding that introducing multiple mutations into the WT protein rendered it a more effective aggregation catalyst for W42Q (Fig. 1D) raised the possibility that the catalytically active conformation of the WT is in some way distinct from the native conformation. Examination of the structure, as well as previous simulations (43, 63), indicated that the only partially unfolded conformation expected to be accessible in WT HγD under physiological conditions is the one where the cores of both domains are intact, but the domain interface is opened. *We hypothesized that this interface-opened conformation was the one active in conformational catalysis.* The free energy of the domain interface has been estimated at just 4 kcal/mol (55), so WT is expected to populate this intermediate, albeit rarely and transiently, under physiological conditions. As described above, PEGylation experiments (Fig. 2 and Fig. S4D) supported interface opening in the CCCC variant; the kinetics of Cys reactivity in this variant supported that this opening was rare. The rarity of this intermediate explains the relatively high concentrations of WT required for catalysis of aggregation: *it is not the native state WT that is catalytically active, but only the minor fraction that adopts the open-interface intermediate conformation at any given time*.

To test the hypothesis of conformational catalysis via the interface, we prepared a series of four constructs expected to have distinct interface-opening propensities and evaluated their ability to catalyze W42Q aggregation. Construct 1 was WT bearing an N-terminal KHHHHHHQ tag sequence; this tag has been shown to interact with the domain linker and part of the C-terminal domain (64), so we reasoned it should populate the open-interface intermediate less frequently than WT due to essentially an auxiliary interface. Construct 2 was the untagged WT protein as used elsewhere in this study. Construct 3 was the H83P variant, which introduces a second Pro residue into the 5-residue domain linker and was expected to increase linker strain and therefore interface opening. Construct 4 was the double-deletion variant Δ85,86, in which two of the five residues that form the domain linker are deleted, which was expected to result in large linker strain. Indeed, purification of the Δ85,86 variant revealed a modest shift toward earlier elution position by size-exclusion chromatography, consistent with a more open structure, and solutions of Δ85,86 became cloudy if kept on ice or even at 4 °C, in contrast to all other crystallin constructs in this study; the cloudiness dissipated completely upon warming the sample to 37 °C or room temperature. The reversible nature of this aggregation process suggests reversible polymerization exclusively at cold temperatures, probably via domain swapping at the opened interface. A fifth “construct” was an equimolar mixture of isolated N- and C-terminal domains. No turbidity at 0 or 4 °C was observed in that mixture.

**Fig. 3:**
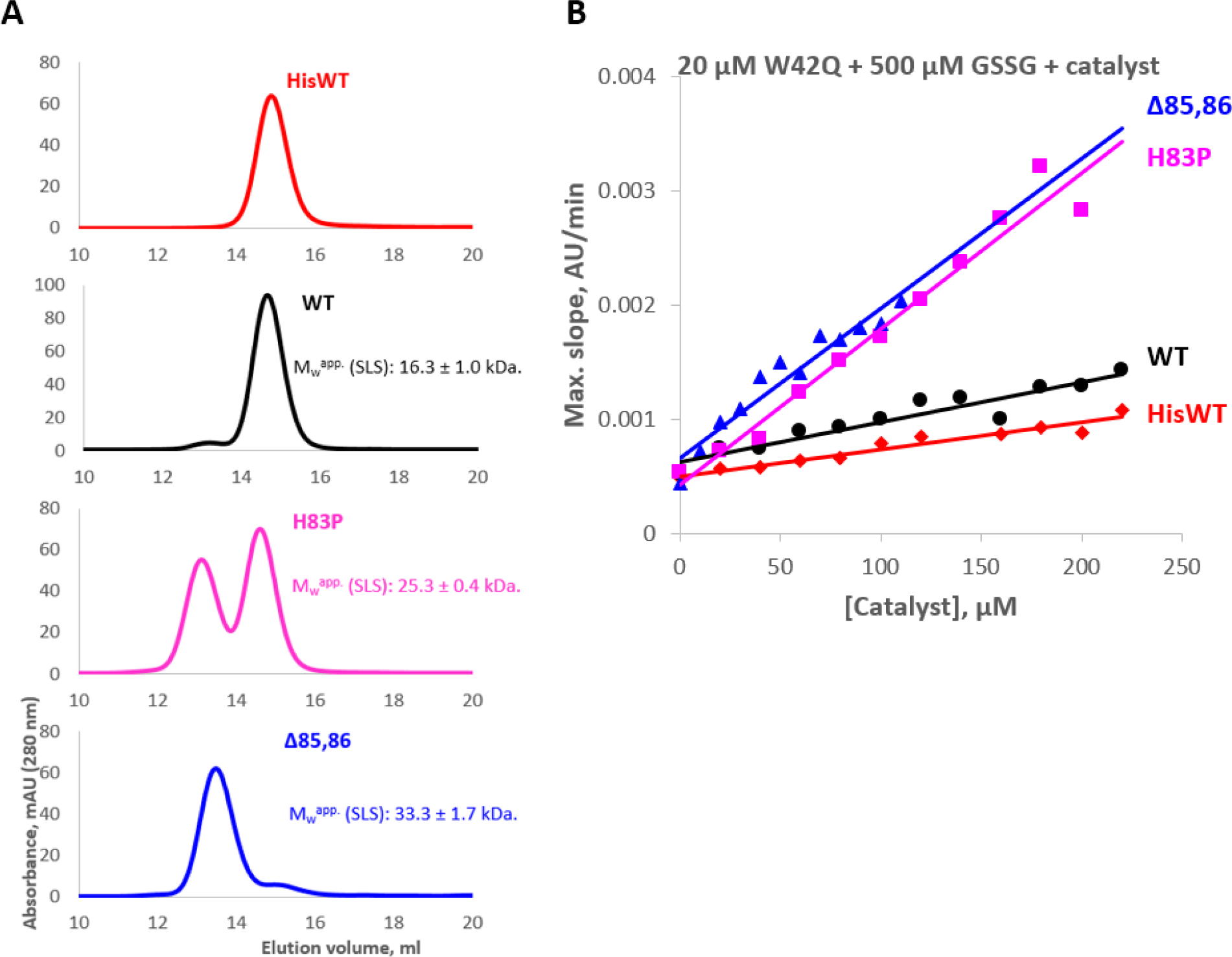
Conformational catalysis is associated with the domain interface. (A) Extent of dimerization by size-exclusion chromatography was in the order of predicted interface-opening propensity. Samples were incubated with reducing agent prior to SEC. Dimerization was confirmed by static light scattering, as indicated by the apparent molecular weights in batch mode. (B) Aggregation-promoting ability of His-WT, WT, H83P, and Δ85,86 had the same rank order as expected accessibility of the domain interface.

As shown in Fig. 3A, the oligomeric state of the four constructs varied in accordance with their expected interface-opening propensities: no detectable dimer in HisWT; an extremely weak dimer peak in WT; a strong one for H83P; and the Δ85,86 variant was predominantly dimeric. Static light scattering experiments in batch mode confirmed the shift to dimeric structures, as indicated. In the case of Δ85,86 the thermostability of both domain cores was lowered (Table 1), consistent with the loss of the interface between them. Surprisingly, such destabilization was not detected in H83P, perhaps because Pro isomerization is relatively rapid at the elevated temperatures used for calorimetry, and the linker strain is thus relieved. Data were analyzed by plotting maximal slope of turbidity increase vs. concentration of catalyst, calculated as in the example of Fig. S5.

The magnitude of conformational catalysis by these four constructs likewise ranked in accordance with their predicted propensity to populate the open-interface intermediate. HisWT was the least active, followed by WT, H83P, and finally Δ85,86 (Fig. 3B). Moreover, a mixture of isolated N- and C-terminal domains had a comparable catalytic activity to the Δ85,86 variant (Fig. 4A). Thus, the WT protein’s catalytic activity was indeed likely related to interface opening.

While both isolated domains produced enhanced turbidity when mixed with the W42Q variant, the reasons for the enhancement were distinct. As shown in Fig. 4B, the isolated N-terminal domain (N-td) co-aggregated with the W42Q variant, whereas the C-terminal domain (C-td) did not. Likewise, the aggregated fraction in the W42Q/Δ85,86 mixture contained ~10% Δ85,86, a minor but significant amount (Fig. S6). It is known that the domain interface in the native HγD structure disproportionately stabilizes the N-terminal domain, both thermodynamically and kinetically (reducing its rate of unfolding) (55, 65). It has been shown that the same non-native disulfide bond (32-SS-41) that is required for aggregation of W42Q also forms in the isolated N-terminal domain (56). Disrupting the stabilizing domain interface thus likely facilitates misfolding and disulfide-trapping of the N-terminal domain even in the absence of any mutations in the core, leading to the possibility of co-aggregation. At the same time, aggregation in mixtures of W42Q and isolated N-td was weaker than in mixtures of W42Q and isolated C-td (Fig. 4A). The observation of weak enhancement and co-aggregation in N-td+W42Q, but strong enhancement and no co-aggregation for C-td+W42Q indicates that the observed interface-mediated conformational catalysis is attributable to the C-terminal side of the interface.

**Table 1:** Thermostability of select HγD variants by differential scanning calorimetry.

| Protein | T<sub>m</sub> 1 (N-td) | T<sub>m</sub> 2 (C-td) |
| --- | --- | --- |
| W42Q <sup>a</sup> | 54.6 ± 0.3 | 71.1 ± 1.7 |
| L5S | 58.3 ± 0.02 | 70.5 ± 0.1 |
| V75D | 60.0 ± 0.01 | 71.0 ± 0.02 |
| WT | 81.2 ± 0.1 | 84.1 ± 0.03 |
| H83P | 81.5 ± 0.1 | 84.2 ± 0.03 |
| $\Delta 85,86$ | 71.6 ± 0.2 | 74.3 ± 0.04 |

Transition midpoints were obtained by fitting the differential scanning calorimetry traces to a double-Gaussian model using the NanoDSC software. Values are reported as mean ± standard deviation of 2-3 technical replicates. <sup>a</sup>Data from reference (43).

To determine whether the catalytic effect on W42Q aggregation was due to specific features of the C-terminal domain interface or to its overall hydrophobicity, we studied aggregation of W42Q in the presence of varying concentrations of bovine serum albumin (BSA). BSA natively has ~28 times more hydrophobic surface area (HSA) per molecule than either of the HγD single domains (and ~60 times more than native HγD WT), as summarized in Table S7. Some acceleration of W42Q aggregation was indeed observed in the presence of BSA (Fig. S8), but the HγD constructs produced much more rapid aggregation than BSA, despite their much lower surface hydrophobicity than BSA. Likewise, HSA for the N-td is higher than for the C-td, yet it is the C-td that is responsible for conformational catalysis. Thus, the observed conformational catalysis is best explained by a specific binding interaction, not merely presence of a hydrophobic surface.

**Fig. 4:**
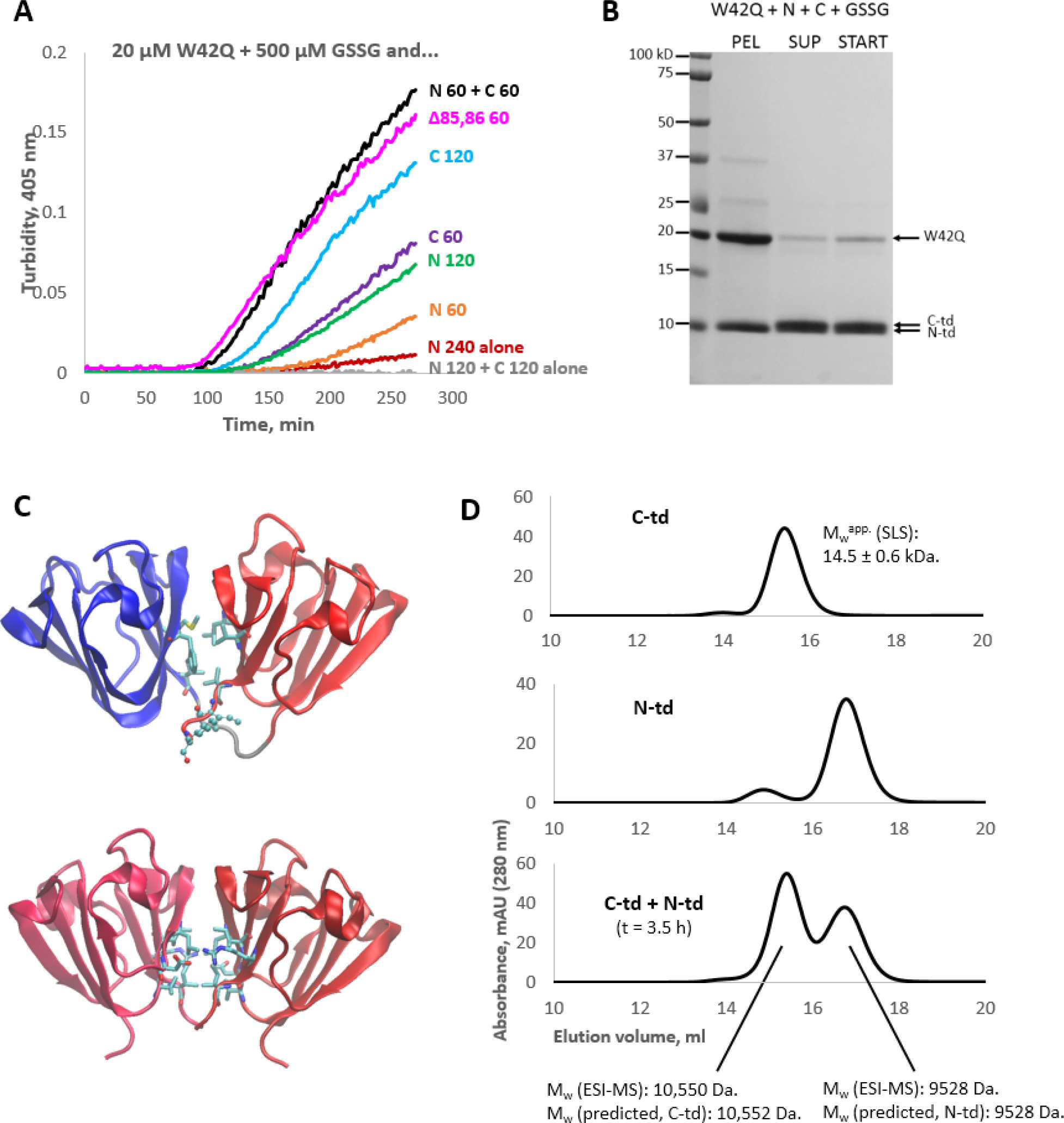
Homodimerization of the C-terminal domain in preference to native N-td:C-td heterodimerization accounts for conformational catalysis. (A) The C-terminal domain promoted aggregation most efficiently, although when N-td and W42Q were both present, aggregation was enhanced to the same level as the Δ85,86 variant. (D) The N-terminal domain co-aggregated with the W42Q variant in approximately stoichiometric amounts, whereas the C-terminal domain showed no co-aggregation. (C) A comparison between the native-state N-td:C-td domain interface of HγD (PDB ID 1HK0) (52) and the interface of the HγS C-td dimer (PDB ID 1HA4) (66). Ile171 and Phe172, in ball-and-stick representation in the HγD structure, would become available as part of a putative C-td-C-td interface. (D) Size-exclusion chromatography of isolated C-td (*top*), isolated N-td (*middle*), and a mixture of N-td and C-td following a 3.5 h incubation at 37 °C in the presence of DTT and ETDA. The mixture experiment did not result in an elution peak containing a complex of N-td and C-td; rather, the two domains eluted separately, N-td as a monomer and C-td as a dimer, as confirmed by electrospray mass spectrometry. The C-td:C-td dimer therefore appeared to outcompete the native-like N-td:C-td pairing.

Investigation of the isolated C-terminal domain provided a structural explanation for the catalytic action of the C-terminal domain interface. It has been shown (66) that the isolated C-terminal domain of human γS-crystallin forms crystallographic dimers with a stable hydrophobic dimer interface composed of the same residues as the domain interface in the native state (Fig. 4C). We found that the HγD C-terminal domain also formed a dimer, eluting close to the full-length WT position by size-exclusion chromatography (Fig. 4D). Dimerization was confirmed by static light scattering in batch mode, as indicated. Furthermore, in mixtures of N-td and C-td, the C-td remained dimeric for hours with no detectable formation of N-td:C-td heterodimers (Fig. 4D and Fig. S9), indicating that in the absence of a linker the C-td:C-td interface is favored over the N-td:C-td one. This could be rationalized by observing that two hydrophobic residues (Ile171 and Phe172) are available for interaction as part of the homodimer, but not in the heterodimer (Fig. 4C).

We propose that transient dimerization of HγD molecules in the open-interface intermediate state via formation of non-native intermolecular C-td:C-td interfaces eliminates stabilization of the N-terminal domain by the native N-td:C-td interface and thus facilitates N-td misfolding. In the case of WT that opening of the interface is too rare and transient, and the N-terminal domain core too stable, to generate detectable misfolding of the N-terminal domain core to form the aggregation-prone 32-SS-41 non-native disulfide. However, when the N-terminal domain core is destabilized by the W42Q or other cataract-associated mutations – or when interface opening is a permanent feature, as in the case of the Δ85,86 variant – the relatively labile N-terminal domain misfolds to populate the aggregation-prone intermediate. In this way, interaction of two HγD molecules in similar intermediate states (open-interface) facilitates their conversion into a distinct and more unfolded intermediate state (hairpin-extruded) and thus accelerates aggregation. This structural model is summarized in Fig. 5A.

**Fig. 5:**
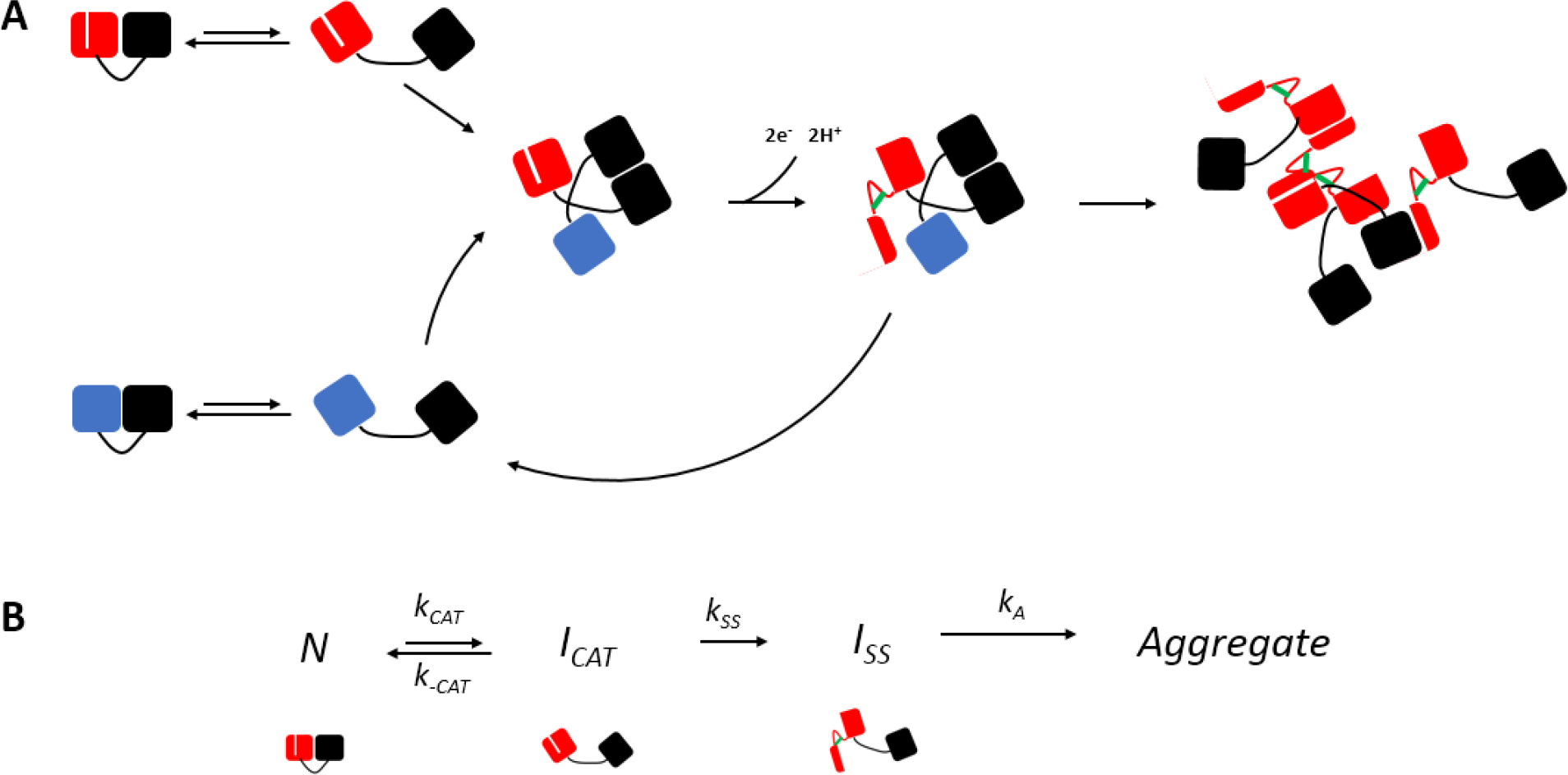
Structural model of conformational catalysis. (A) The catalytic interaction depends on a complex between cognate domains of two molecules (black), which deprives their partner domains (red, blue) of the added kinetic stability contributed by the native domain interface. If this partner domain also bears a destabilizing modification (white stripe), misfolding becomes likely. In the case of HγD, the misfolded conformer is locked by a non-native disulfide bond (green), while the wild-type intermediate is free to dissociate without further misfolding in a reaction driven by re-formation of the native domain interface and the high native solubility of WT. The aggregate model depicts the three recognized interaction modes between the detached N-terminal hairpin of HγD W42Q and sites on other copies of the protein, based on (43). (B) The structural model gives rise to a kinetic scheme for “inverse prion” aggregation involving two distinct partially unfolded intermediates, ***I***_***CAT***_ (interface-opened) and ***I***_***SS***_ (hairpin-extruded due to the 32-SS-41 disulfide), the earlier of which promotes formation of the latter. The equilibrium in the first step favors the native state (***I***_***CAT***_ is rare), and the conversion of ***I***_***CAT***_ to ***I***_***CAT***_ occurs only in the destabilized mutants and is rate-limiting for the aggregation reaction.

The structural model of conformational catalysis via interface stealing can be formalized in the kinetic scheme depicted in Fig. 5B. Both the WT protein and the aggregation-prone mutants (***MUT***) fluctuate between the native (***N***) state, which we define here as any state that is neither aggregation prone nor likely to catalyze aggregation, and the ***I***_***CAT***_ state, an early conformational intermediate with an opened domain interface. ***I***_***CAT***_ is capable of catalyzing aggregation but is not itself aggregation prone. Its concentration is determined by ***K***_***CAT***_ = ***k***_***CAT***_/***k***_-***CAT***_, and it is thus proportional to the starting concentration of protein – in the case of ***MUT***, at early times, and in the case of WT, always. Only ***MUT*** can proceed from ***I***_***CAT***_ to ***I***_***SS***_, a partially misfolded conformation in which the N-terminal β-hairpin is extruded and precluded from re-annealing to the native position due to formation of the intramolecular 32-SS-41 disulfide bond (43).

***I***_***SS***_ is the aggregation precursor, and the conversion from ***I***_***CAT***_ to ***I***_***SS***_ is the rate-limiting step for the aggregation process, at least during the time when the rate of aggregation is maximal. Prior experimental data support this result. ***I***_***SS***_ in W42Q forms at temperatures where the requisite unfolding is undetectably rare in the absence of a non-native disulfide (43, 57), whereas the domain interface has a relatively low ΔG of unfolding (4 kcal/mol) (55) and has been shown to open rapidly in both Monte-Carlo (63) and molecular dynamics (62) simulations. The observation that ***I***_***SS***_ partitions quantitatively to the aggregated state with none detectable in the soluble fraction at 37°C and pH 7 (44) indicates that ***k***_***A***_ is not rate-limiting.

We have previously shown an approximately linear correlation between solution turbidity and amount of aggregated W42Q at early times in the aggregation reaction (57). The structural model and kinetic scheme in Fig. 5 make clear predictions about the concentration dependence of aggregation rates at such times:

First, since the rate-limiting step, formation of ***I***_***SS***_, is bimolecular in ***I***_***CAT***_, its rate should be quadratic in ***[I_CAT_]***. In turn, ***[I***_***CAT***_] = ***K***_***CAT***_***[N]***, which is proportional to the starting concentration of the mutant at early times. Then, the maximal aggregation rate for ***MUT*** alone should vary as the square of its starting concentration. This is indeed observed (Fig. 6B).

Second, since conformational catalysis by WT implies formation of a mixed dimer of ***I***_***CAT***_***(WT)*** and ***I***_***CAT***_***(MUT)***, the concentration dependence of maximal rate on [WT] is expected to be linear. This is also observed in every case (Fig. 6D), although the linearity is imperfect for W42Q. A likely explanation is that the W42Q mutation destabilizes the domain interface along with the N-td core, resulting in strong self-catalysis. At high [WT] heterodimers of ***I***_***CAT***_ predominate, and the contribution from self-catalysis is reduced. Depletion of W42Q may also play a role at the low concentrations (20 μM) used in this experiment.

Third, since conformational catalysis requires dimerization of the C-terminal domains, aggregation of isolated wild-type N-terminal domain is expected to proceed without self-catalysis. This should yield a linear concentration dependence of maximal slope on starting concentration, in contrast to the quadratic dependence for the full-length variants. A linear concentration dependence of maximal rate is indeed observed (Fig. S10A).

Fourth, since conformational catalysis proceeds via interface stealing, and heterodimers in mixtures of the isolated domains do not form (Fig. 4D), a physical linkage between N-td and C-td must be required. Hence, aggregation in an equimolar mixture of isolated N-td and C-td is expected to still have a linear concentration dependence, as opposed to quadratic dependence if a direct misfolding-promoting N-td/C-td interaction were involved. Linear dependence is indeed observed (Fig. S10B), confirming that conformational catalysis proceeds via interface stealing in the two-domain protein.

In the following section we present a more comprehensive and quantitative kinetic model of the aggregation process and also show that this model does not depend on any particular relationship between solution turbidity and amount of aggregated protein.

### Kinetic model of conformationally catalyzed HγD aggregation

The central assumption of this model is that the formation of the I_SS_ state and the subsequent aggregation of I_SS_ monomers are independent, irreversible processes. This independence means that the presence of aggregates does not affect the kinetics of misfolding, and misfolding only affects the aggregation process through the time-dependent population of I_SS_ monomers. We shall show that, given this assumption and the kinetic scheme of Fig. 5B, we can use turbidity data to study the kinetics of catalyzed misfolding.

To make this connection, we first write a differential equation for the population of protein in the state I_SS_, denoting the total flux of protein into this state by *J*_SS_,

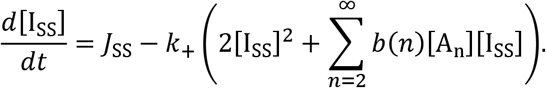

The second term on the right-hand side represents the two ways by which an I_SS_ monomer can aggregate: by forming a new aggregate with another disulfide-locked monomer, or by attaching to an existing aggregate *A* with *n* monomers and concentration [A_*n*_]. The aggregation rate constant, *K*_+_, is the same for all aggregate sizes, but larger aggregates may expose multiple binding sites *b(n)*. If we assume for the moment that *J*_SS_ is constant (i.e., depletion of N due to aggregation is negligible), then we can eliminate the dependence on both *J*_SS_ and *k*_+_ by rewriting the equation in terms of a scaled time 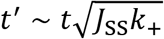 and scaled concentrations 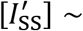 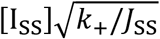 and 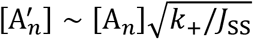 (see **SI text**). This transformation implies that the distribution of aggregate sizes 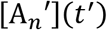 is independent of *J*_SS_. In reality, *J*_SS_ is not exactly constant due to the depletion of native-state protein. However, deviations from the ideal scaling relationship depend on the details of the misfolding kinetics, as we describe for each distinct case below.

We can use our scaling prediction to analyze the turbidity curves without knowing the size, shape, or light-scattering properties of the aggregates (67). Since the turbidity, τ, is linear in the concentration of aggregate particles, application of our scaling prediction implies that the turbidity traces, in the absence of substantial depletion, should collapse onto a single master curve, 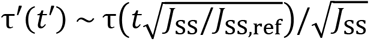. In practice, we choose the reference state to be the turbidity trace at the highest protein concentration. Fig. S11 shows that, for all mutant—catalysis combinations, the highest concentration turbidity traces do indeed collapse onto a master curve. We can verify this scaling prediction quantitatively by calculating the time at which the turbidity trace crosses a fixed threshold value of the scaled turbidity τ′ (Fig. 6A,C). Alternatively, we can extract the dependence of *J*_SS_ on the protein concentration by analyzing the slope of the turbidity trace at a fixed scaled time 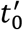 on the master curve (Fig. 6B,D). Assuming negligible depletion, this slope is proportional to *J*_SS_,

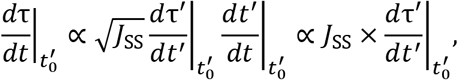

regardless of the shape of τ′(*t*′). In summary, the general assumptions of irreversibility and catalysis—aggregation independence as outlined above, taken together with the empirically verified scaled data collapse, make it possible to study the mechanism of *I*_SS_ formation by analyzing the concentration-dependent turbidity curves. We now consider the distinct cases of self-catalysis and heterogeneous catalysis:

#### Case I: Self-catalysis

In this case, the rate-limiting step is facilitated by an interaction between aggregation-prone molecules in the catalytic intermediate state. Hence,

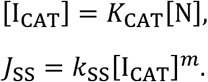

The equilibrium constant *K*_CAT_ = *k*_CAT_/*k*_−CAT_ is variant-specific. The simplest possibility is that the catalytic complex consists of only two I_CAT_ molecules, so that *m* = 2 and the dependence of *d*τ/*dt* on [N] is quadratic. This concentration-dependence is in excellent agreement with the experimental data shown in Fig. 6B, which also demonstrates that a linear fit fails.

Interestingly, for this form of catalysis, the effect of depletion on the scaling transformation does not depend on the initial concentration of native-state protein (see **SI text**). In agreement with this prediction, we observe no systematic deviations from the predicted time scaling shown in Figure 6A or the master turbidity curve (Fig. S11) as we vary the concentration of each mutant.

By contrast, the rate data for the isolated N-terminal domain, where no catalysis is expected due to the key role of the C-terminal domain in conformational catalysis, do not exhibit a quadratic dependence on concentration (Fig. S12C). The line of best fit has a non-zero intercept, possibly indicating a minimum concentration below which aggregation does not occur. The times at turbidity threshold for the isolated N-td also deviate from the predicted scaling (Fig. S12B), which could indicate a strong depletion effect or a kinetic scheme for aggregate nucleation distinct from that of the full-length variants; this will be a subject of future research. Nevertheless, the linearity of the dependence of maximal slope on [N] strongly suggests this reaction is not self-catalyzed, in agreement with the prediction from the structural model (Fig. 5A).

#### Case II: Heterogeneous catalysis

This case differs from Case I in only one key respect: Catalysis can be carried out not only by the I_CAT_ state of the mutant, but also by the equivalent conformation of a non-aggregation-prone variant. Using the simple model of a bi-molecular catalytic complex, which was validated in Case I, we have

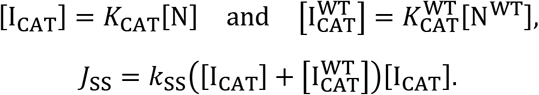

Note that since the non-aggregation-prone variant (which we label WT for simplicity) does not proceed to the I_SS_ conformational state, 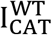 is determined by an equilibrium process. When the concentration of the mutant protein is fixed and depletion is negligible, the flux of misfolded protein depends linearly on the heterogeneous catalyst concentration,

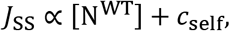

which is in excellent agreement with the measurements of *d*τ/*dt* shown in Fig. 6D. The mutant-specific constant *c*_self_ accounts for the modest but non-negligible contribution of self-catalysis to the production of *I*_SS_ monomers under these conditions.

Depletion due to heterogeneous catalysis leads to deviations from the scaling predictions that increase as the WT concentration is increased or the rate of catalysis is increased (see SI text). Such systematic deviations from the ideal time scaling can be seen at high WT concentrations in Fig. 6C,D. Because depletion does not affect the concentration scaling of self-catalyzed aggregation, these deviations from ideal scaling provide an additional, independent test of heterogeneous versus self-catalysis. Another likely contributing factor is reduced W42Q self-catalysis at high [WT], as discussed above.

**Figure 6:**
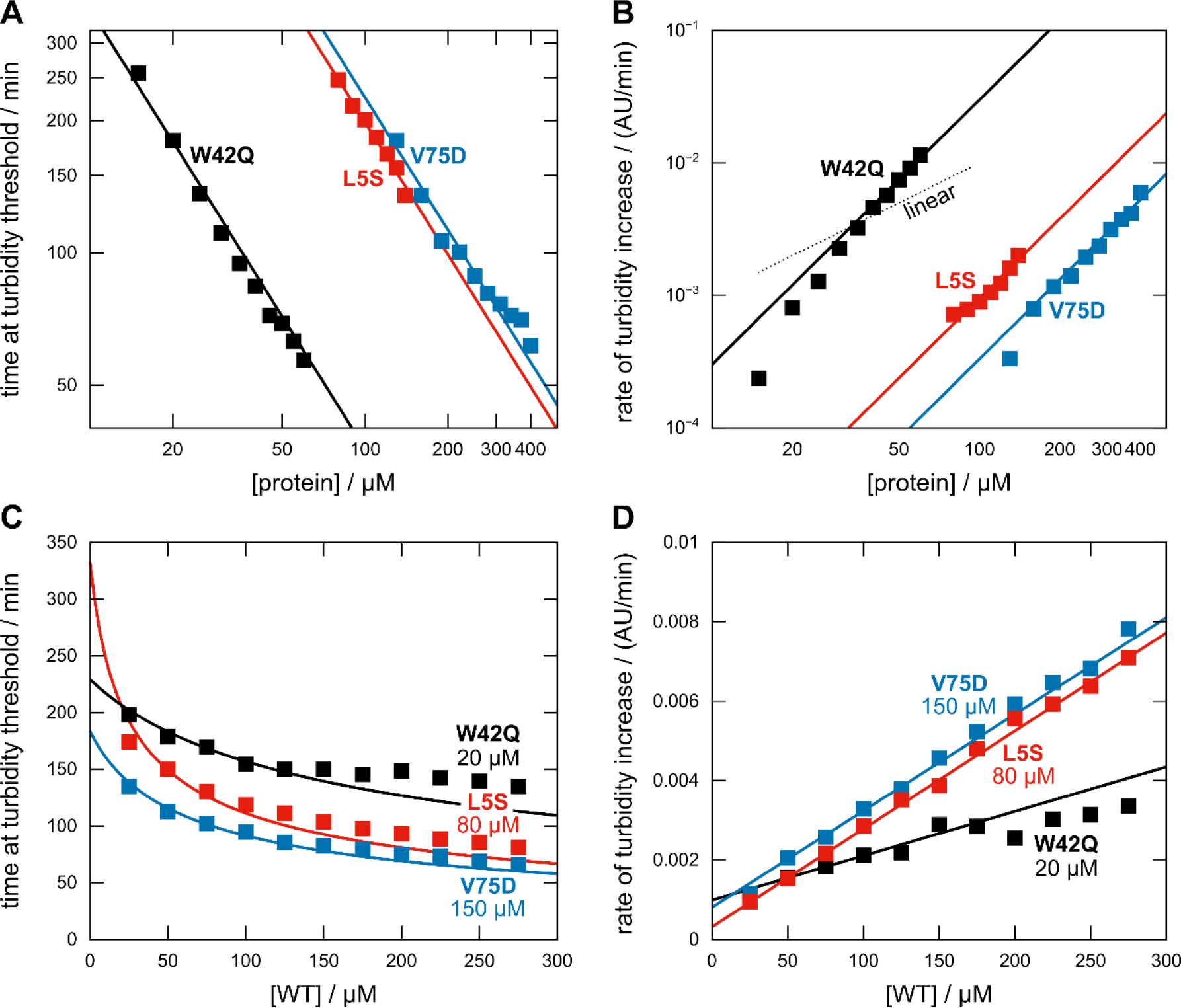
Aggregation times and rates depend on the concentrations of aggregation-prone W42Q, V75D, and L5S in accordance with scaling predictions. We calculated the times when the turbidity traces cross a fixed scaled turbidity threshold (panels A,C) and the slope of the turbidity traces at a fixed scaled time (panels B,D; see Methods and SI Figure 1). In the case of self-catalysis (panels A,B), the times and rates are proportional to [N]^−1^ and [N]^2^, respectively; these fits appear as straight solid lines on the log—log plots. For heterogeneous catalysis (panels C,D), the rates are fit to the linear function *J*_SS_ ∼ [N^WT^] + *c*_self_ using mutant-specific constants *c*_self_ = 88, 33, and 13 μM for W42Q, V75D, and L5S, respectively. Systematic deviations from the ideal time scaling 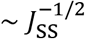 are observed at high WT concentrations (panel C), likely due to depletion. WT appears to be a stronger catalyst of L5S and V75D aggregation (panel D), but this distinction is likely due to the higher concentrations of these mutants relative to W42Q in these experiments.

## Discussion

In previous work we have shown that both misfolding and formation of a nonnative internal disulfide are required for the aggregation process in cataract-associated variants of HγD (43, 57). This disulfide locks a specific aggregation-prone structure, and oxidation of the same two Cys residues in aged patients without any mutations was found to correlate with cataract severity (35). This convergence supports the growing understanding that mutational and post-translational damage lead to misfolding via highly similar mechanisms (45, 58). We have also shown the surprising ability of HγD, and likely other γ-crystallins, to dynamically exchange disulfide bonds, potentially serving as a redox buffer (44). The “hot potato” mechanism proposed in (44) asserts that WT HγD has an oxidoreductase activity by oxidizing other HγD molecules in a relay-like manner until a mutationally or post-translationally destabilized HγD molecule gets oxidized with subsequent formation of the 32-SS-41 disulfide causing irreversible aggregation and elimination of the disulfide from solution. In this way, oxidized WT promotes aggregation of W42Q or other destabilized variants. This mechanism, too, found support in lens proteomic studies (37, 42).

We have now discovered a new mechanism by which WT HγD can promote aggregation of aggregation prone variants independent of any redox activity. Even oxidoreductase-inert versions of WT HγD (such as CCCC) dramatically increased aggregation kinetics and conformational dynamics of cataract-associated W42Q HγD. Aggregation of other cataract-associated variants (V75D and L5S) was accelerated even more. The mechanism at work is *conformational catalysis*. In the conformational catalysis mechanism, partial unfolding of one HγD molecule catalyzes misfolding of another. The observation of strong C-td:C-td association, combined with prior findings of significant destabilization of the isolated N-td, led us to propose that conformational catalysis in HγD proceeds by interface stealing. Our structural model pointed to formation of a key bimolecular complex between wild-type and mutant, or mutant and mutant, molecules that serves as a catalyst of conformational conversion of the mutant to the aggregation-competent form. This motivated a kinetic model based on the assumption that conformational catalysis and aggregate assembly are fully independent processes. The power-law relationships between protein concentration and rate of turbidity rise and time before the turbidity threshold (“lag time”) derived from the model are in excellent agreement with the experimental data (Fig. 6).

It’s important to note that the observed aggregation pathway does not depend on any particular mutation: oxidative aggregation under physiologically relevant conditions occurs not only for W42Q, but also for V75D, L5S, and even fully wild-type N-terminal domain, as long as the interface with the C-terminal domain has been disrupted. Disruption of the domain interface need not even require drastic damage, since it has been shown that one or two deamidations can be enough (54, 68), and deamidation in the lens is common (24). These findings indicate that a variety of perturbations to diverse areas of the protein’s structure all converge on similar partially unfolded aggregation-prone intermediates and pathways of assembly.

The peculiar structural mechanism of conformational catalysis via interface stealing by the catalyst protein from the aggregation-prone variant (Fig. 5A) is rooted in the evolutionary history of the γ-crystallins, whose two domains arose by gene duplication and fusion (69). The domains are structurally identical, but their sequences have diverged (37% identity). Structural similarity promotes propensity of proteins to interact (70, 71). Further, it was shown that for structurally similar domains, sequence divergence weakens their interaction propensity (72), consistent with our observation that homodimerization of C-td is more favorable than C-td:Ntd heterodimerization (Fig. 4D). The C-terminal domain of human γS-crystallin can form dimers (66), and in that protein, too, loss of the native domain interface greatly accelerates unfolding of the Cys-rich N-terminal domain (55, 65). Dimerization of the N-terminal domains via disulfides (73) may serve a protective function in this scenario. We expect that conformational catalysis like that presented here for HγD may also operate in HγS, other members of the γ-crystallin family, and likely many other multidomain proteins.

A direct test for interface stealing *in vivo* is not possible with current technology. However, isolated C-td was reported to arise in aged lenses by spontaneous domain linker cleavage, though no N-td was recovered (74), which indirectly supports the physiological relevance and consequences of interface stealing.

In the WT protein, interface stealing is likely not prevalent for entropic reasons – due to the high effective concentration of the two domains tethered by only a 5-residue linker. Furthermore, it has been shown that protein stability contributes to the strength of interaction with other proteins (75, 76). Interface stealing could be an evolutionary side effect of domain duplication and fusion. When the N-td is destabilized, as in W42Q, V75D, and L5S, stability of the N-td:C-td interface is likely also weakened, making the intermolecular C-td:Ctd interface relatively more favorable. This is especially true for mutations at Trp42, which disrupt both the N-td core and the domain interface (62). This double disruption explains why W42Q is much more aggregation-prone than L5S or V75D despite relatively small differences in thermostability (Table 1) – C-td:C-td dimerization is especially favorable in W42Q relative to N-td:C-td, making it a potent self-catalyst. The L5S and V75D mutations are further from the domain interface, so their rates of self-catalyzed aggregation are lower, and aggregation is accelerated by the WT protein to a relatively greater extent. Indeed, the efficiency of self-catalysis relative to catalysis by WT corresponds to the mutation’s proximity to the domain interface: W42Q > V75D > L5S (Fig. S11).

The phenomenon of conformational catalysis is conceptually the opposite of the by now well-known case of conformational templating (as in prions). In the templating case, protein-protein interactions shift one protein molecule into the same conformation as another (77–79). In the present case, partially unfolded WT protein catalyzes the mutant’s conformational conversion to a state that the WT itself does not populate. “Inverse prion” interactions of this type have not been reported to our knowledge previously. Yet, they are likely to play a role in determining the properties of highly concentrated protein solutions. Conformational catalysis is expected to cause conformational heterogeneity to arise more rapidly at high protein concentrations than at low concentrations. This may promote non-native aggregation, but may also inhibit deleterious native-state aggregation, such as crystallization, which is an important evolutionary constraint in the lens (25, 52, 80). Future research should examine whether any defensive mechanisms have evolved to mitigate the aggregation-promoting effect of conformational catalysis.

The general requirements for conformational catalysis to exist are relatively few. The mechanism presented here requires a multidomain protein where the stability of one domain is derived in part from its interface with another, and where interface stealing is possible. Prime candidates are proteins whose interacting domains arose by duplication and fusion. However, interface stealing could also occur between otherwise unrelated proteins that are homologous for one domain. Besides interface stealing, classical domain swapping could also lead to a similar effect, as long as geometric or other constraints permit only a single swap, leaving one of the two destabilized domains unpaired. More generally, whenever a transient interaction between two protein molecules in a non-native conformational state further alters the conformation of one of them, conformational catalysis may be said to occur. Future research should investigate how significant a role conformational catalysis, and interface stealing in particular, may play in the stability, function, and aggregation of multidomain proteins.

## Methods

### Protein expression and purification

Expression and purification were carried out as described in (44). Briefly, untagged HγD constructs were expressed from the pET16b vector in BL-21 RIL cells in SuperBroth medium (Teknova). The cells were grown to late-log or early-stationary phase to obtain the highest yields under our expression conditions. After lysis by sonication, the proteins were purified by ammonium sulfate fractionation (collecting the fraction between 30% and 50% ammonium sulfate), followed by ion exchange on a Q-sepharose column (GE Life Science) and size exclusion on a Superdex75 26×600 column (GE Life Science) in the standard sample buffer, 10 mM ammonium acetate with 50 mM sodium chloride, pH 7. The three mutants whose aggregation traces were used in Figure 3 were treated with a reducing agent (1 mM dithiothreitol) for 2 h at room temperature prior to size exclusion.

### Solution turbidity assays and fitting

Turbidity assays were carried out as described in (44). All samples contained 1 mM EDTA to inhibit any trace metal-induced aggregation as seen in (81–83). Sample volume was 100 μl in half-area polypropylene plates (Greiner Bio-One), resulting in a path length of ∼0.4 cm at the meniscus. Readings were taken in 96-well format on the PowerWave HT plate reader (BioTek). Maximum rate of turbidity was defined as the slope of the steepest tangent, which occurred early in the turbidity trace. Absolute threshold values of turbidity were determined empirically for each mutant; they were 0.02 for W42Q; 0.02 for V75D; and 0.005 for L5S, since the aggregation of that variant appeared biphasic (see Fig. S3). Linear regression fits were applied to the 10 time points starting 1, 2, 3, …, 20 points post-threshold. The resulting slopes were averaged. Scaled turbidity thresholds were set as indicated in Fig. S11, yielding essentially identical results. Fits to the maximum rate vs. concentration and lag time vs. concentration data used linear regression with the equations described in the kinetic model section.

### PEGylation assays

Cys-accessibility PEGylations were carried out as follows. W42Q (20 μM) was incubated with or without the C18T/C78A/C108S/C110S variant (100 μM) and PEG(5000)-maleimide (500 μM) (Sigma) in standard sample buffer at 37 °C in the absence of any oxidizing agent. 5-μl samples were taken at the time points indicated (Fig. 2 and Fig. S4) and mixed with 15 μl of denaturing buffer containing 5 μl 4x Tris/SDS NuPAGE gel-loading buffer (Thermo Fisher), 10 mM tricarboxyethyl phosphine (TCEP) as the reducing agent, and 1 mM L-Cysteine as the quencher of PEG-maleimide. The denaturing buffer was prepared the day of the experiment and allowed to equilibrate for 1 h prior to use to ensure all L-Cysteine was in its reduced form. Three or four replicate samples were used in each case. Cys-counting PEGylations were carried out as described in (44), in the presence of reducing agent (1 mM TCEP). Minor bands at +4PEG in Figure 1E indicate that the reduction of internal disulfide bonds in W42Q was still not 100% complete, possibly because some bonds remained buried during the reductive treatment.

### Analytical size-exclusion chromatography

Samples at 10 μM concentration in standard sample buffer were incubated at 37 °C with 1 mM DTT and 1 mM EDTA for between 2 and 3.5 h. In the case of the N-td+C-td mixture the concentration was 10 μΜ of each. 100 μl of each sample was injected on the Superdex 75 10×300 analytical column (GE Life Science) in standard sample buffer, and peak fractions were collected manually.

### Differential scanning calorimetry

A nanoDSC robotic calorimeter (TA Instruments) was used, following the procedure in (43). Samples were buffer-exchanged by dialysis to 20 mM sodium phosphates, 50 mM NaCl buffer. Reducing agent (1 mM TCEP) was added prior to the experiments.

### Mass spectrometry

MALDI-TOF mass spectrometry was carried out following the protocol of (84) using a stainless steel target plate (Bruker) and the Bruker Ultraflextreme mass spectrometer. Pelleted samples were partially resolubilized with 0.1 M pH 5 ammonium acetate buffer. Samples were desalted using Pierce C18 tips (Thermo Fisher) and eluted in 80% acetonitrile with 0.1% trifluoroacetic acid. For electrospray mass spectrometry, samples from the size-exclusion were desalted using Pierce C18 tips (Thermo Fisher), eluted in 80% acetonitrile with 1% formic acid, and analyzed by LC/MS on an Agilent 6210 electrospray mass spectrometer, followed by maximum entropy deconvolution with 1 Da step and boundaries of 7,000-25,000 Da.

### Static light scattering

Samples at concentrations of 40 μM or (for C-td) 100 μM in standard sample buffer were treated with 1 mM DTT for ~1 h at room temperature to remove any possible C110-C110 dimers. They were then analyzed in 50-100 μl volume on the NanoStar dynamic light scattering instrument (Wyatt) in static light scattering mode, using disposable cuvettes from the manufacturer. The autocorrelation functions were deconvoluted using the Wyatt Dynamics software with the following restraints: 1.5-200 μs correlation times and 1.0-4.0 nm hydrodynamic radii. At least 20 measurements were averaged for each sample.

## Acknowledgements

This work was supported by National Institutes of Health grants GM111955, GM126651, GM068670, and GM124044.

## Supplementary Information

**Fig. S1:**
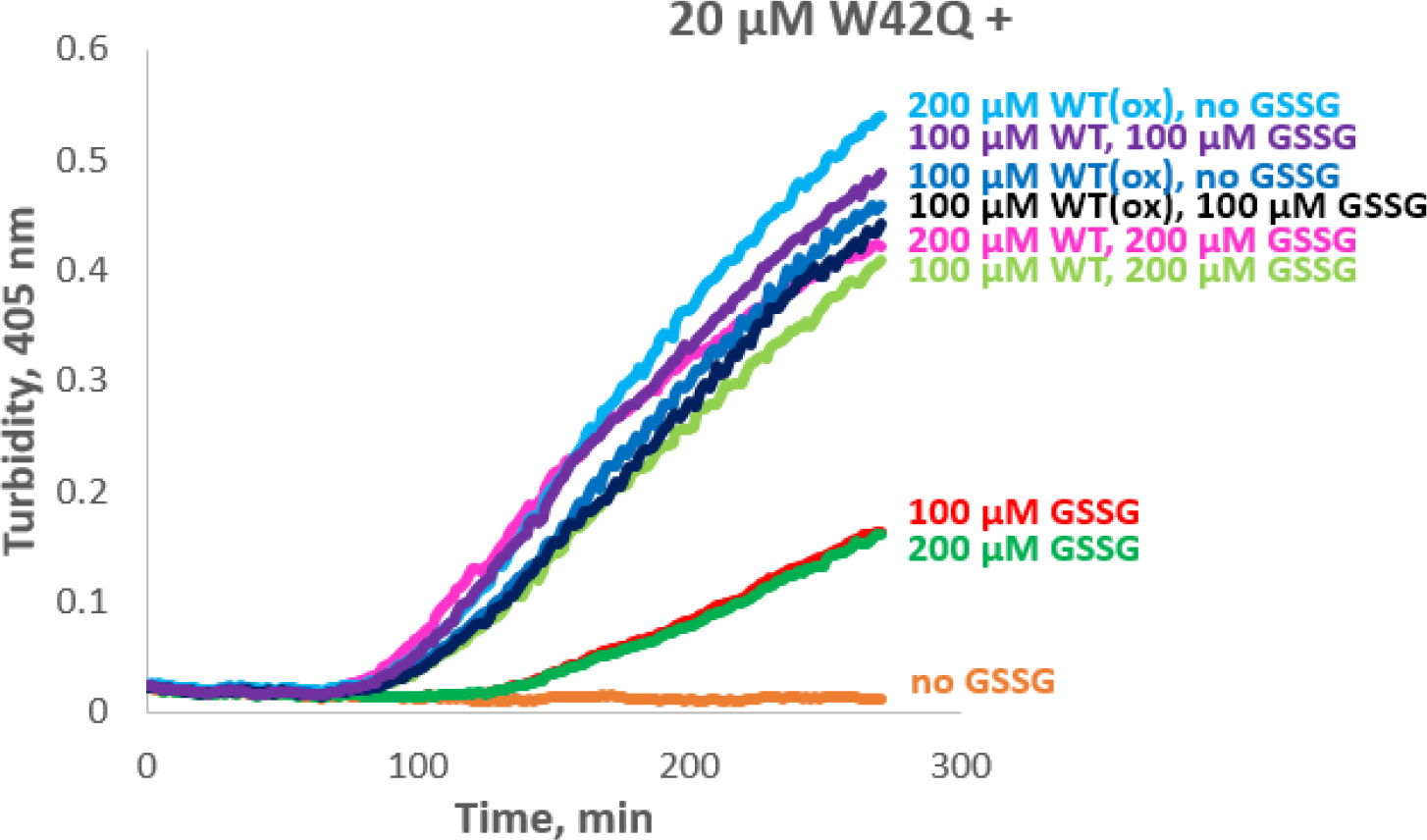
Oxidation status of WT does not affect its ability to catalyze W42Q aggregation when excess GSSG is present. We have previously shown (ref. 42) that in the absence of any oxidation WT does not promote aggregation of W42Q, and that this aggregation requires non-native disulfide formation in W42Q (ref. 41). GSSG alone promoted W42Q aggregation, and so did oxidized WT, both consistent with these prior reports. However, the effect of GSSG + WT, or the WT protein alone if oxidized, was far greater than that of GSSG alone at the same concentration. Indeed, oxidation status of WT made no difference if excess GSSG was present in the sample.

**Fig. S2:**
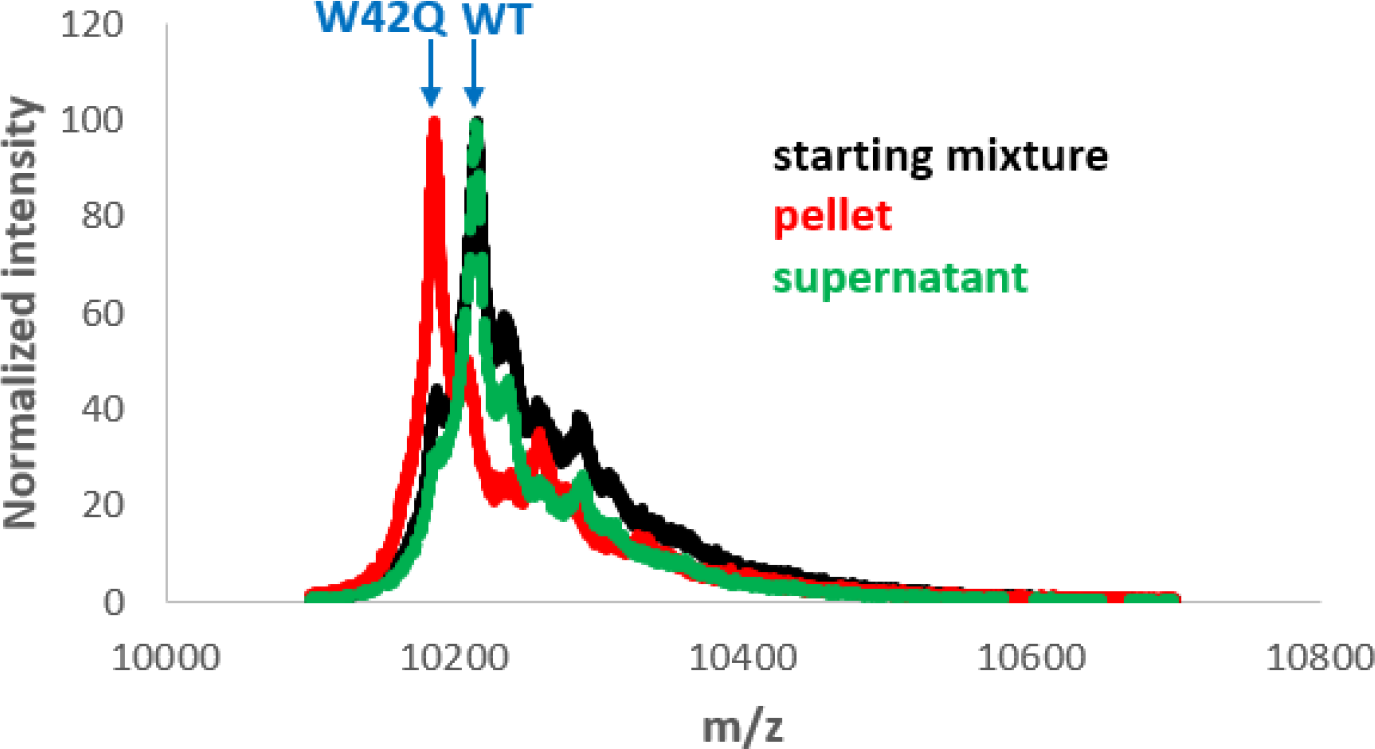
Aggregates formed in the WT + W42Q + GSSG mixture contain only W42Q. MALDI-TOF spectral region containing the z = 2 ion confirmed that the starting mixture (*black*) contained both W42Q and WT, with an excess of the latter; the pelletable fraction contained only W42Q (*red*); and the remaining supernatant (*green*) contained the same amount of WT, but less W42Q, compared to the starting mixture.

**Fig. S3:**
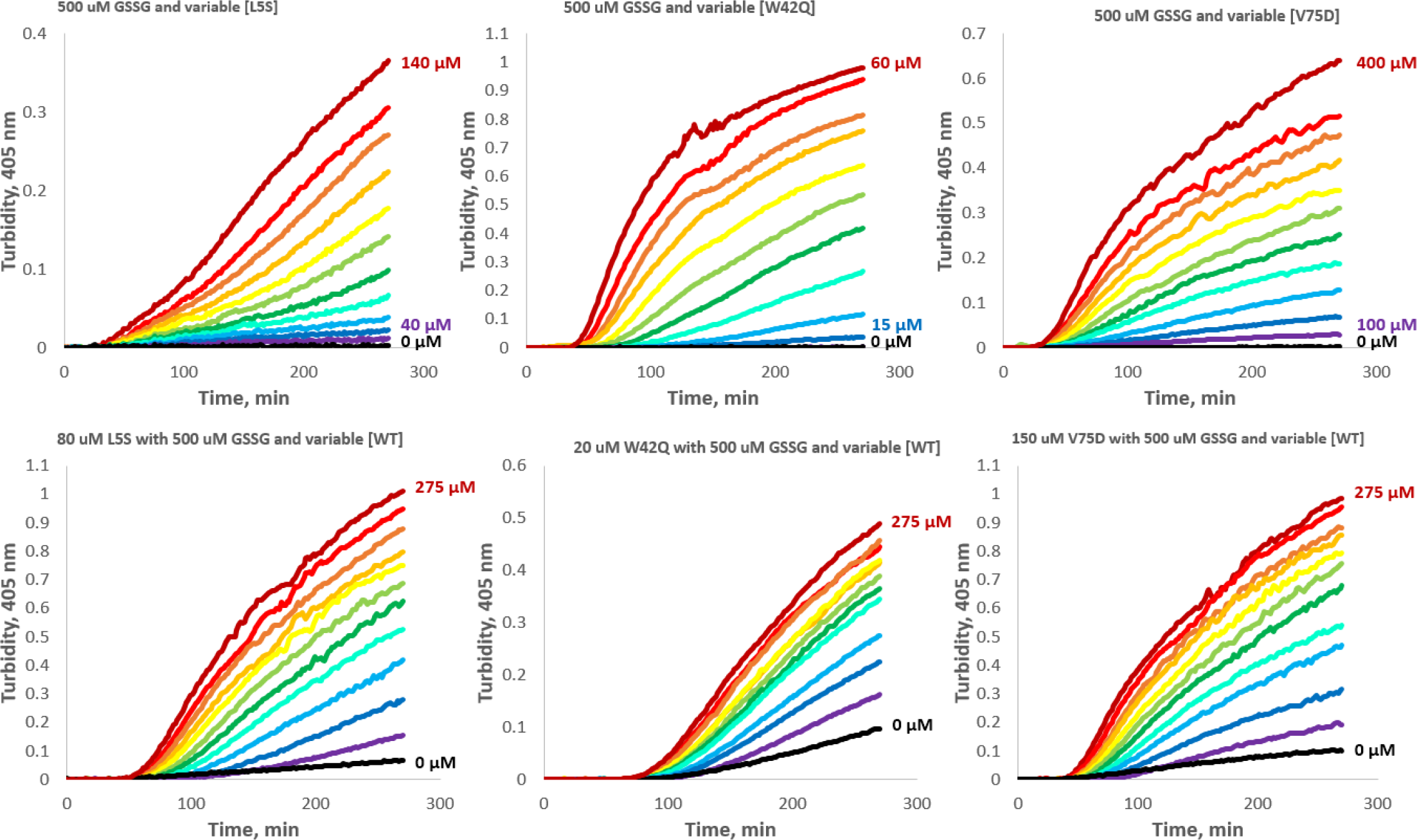
Turbidity traces for homogeneous and heterogeneous (WT-catalyzed) aggregation of the L5S, W42Q, and V75D variants. Concentration of the variable protein increases from blue to red, with the lowest and highest concentrations indicated. Top row: Aggregation of L5S was assayed at concentrations from 40 to 140 μM, with a step of 10 μM; W42Q was assayed from 10 to 60 μM, with a step of 5 μM; V75D was assayed from 100 to 400 μM, with a step of 30 μM. *Bottom row*: Variable amounts of WT protein were added to a fixed amount of each mutant. [WT] was varied from 0 to 275 μM, with a step of 25 μM.

**Fig. S4:**
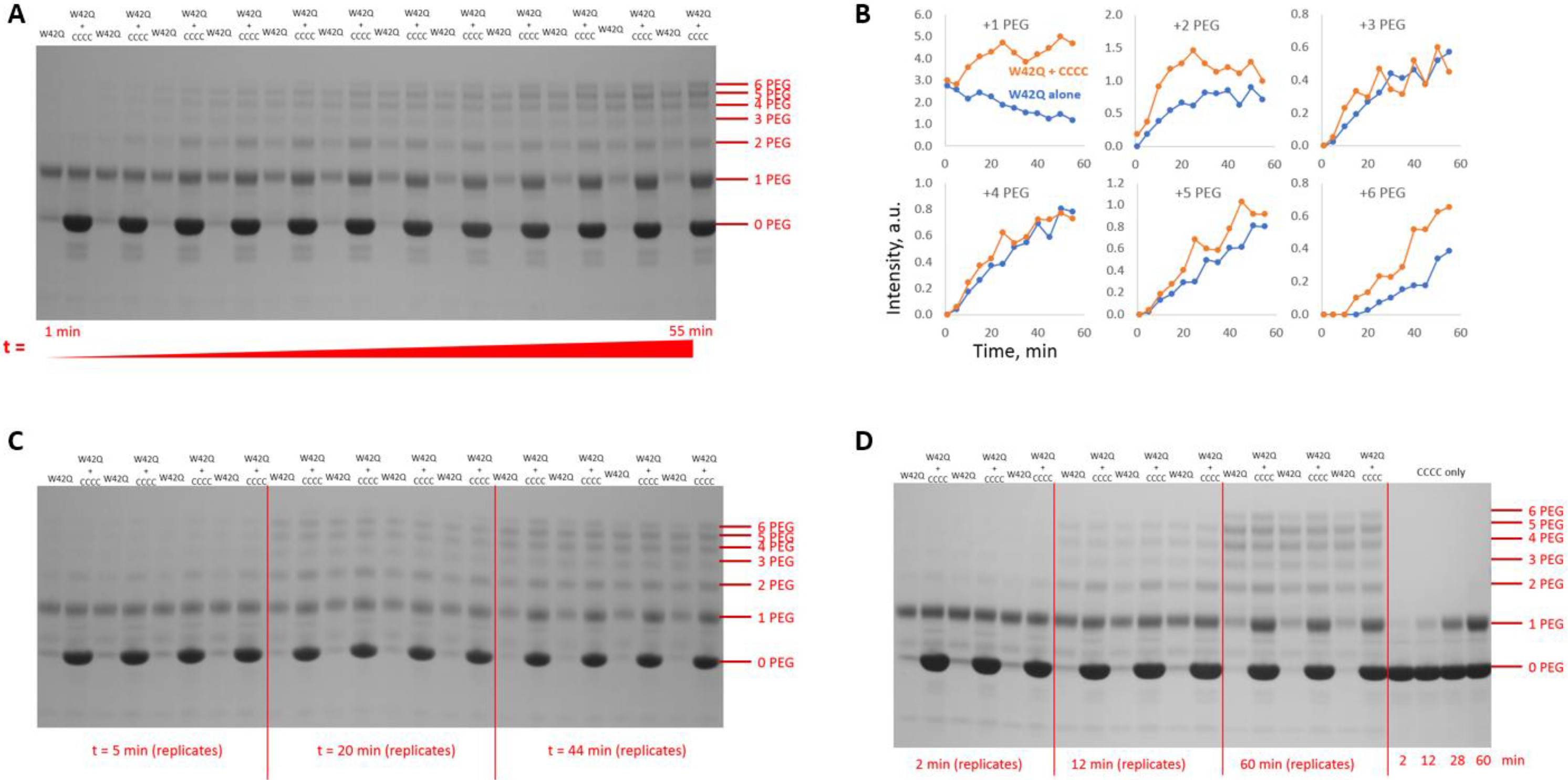
Solvent accessibility of buried Cys residues in the W42Q N-terminal domain increases in the presence of WT analog. (A, C, D) The W42Q mutant sample was incubated with the WT-like quadruple Cys variant C18T/C78A/C108S/C110S and PEG-maleimide, and samples taken at indicated time points were analyzed by SDS-PAGE with subsequent quantitation by gel densitometry using GelAnalyzer 2010 software (B). Quantitation of the gels shown in (C) and (D) is presented in Fig. 2.

**Fig. S5:**
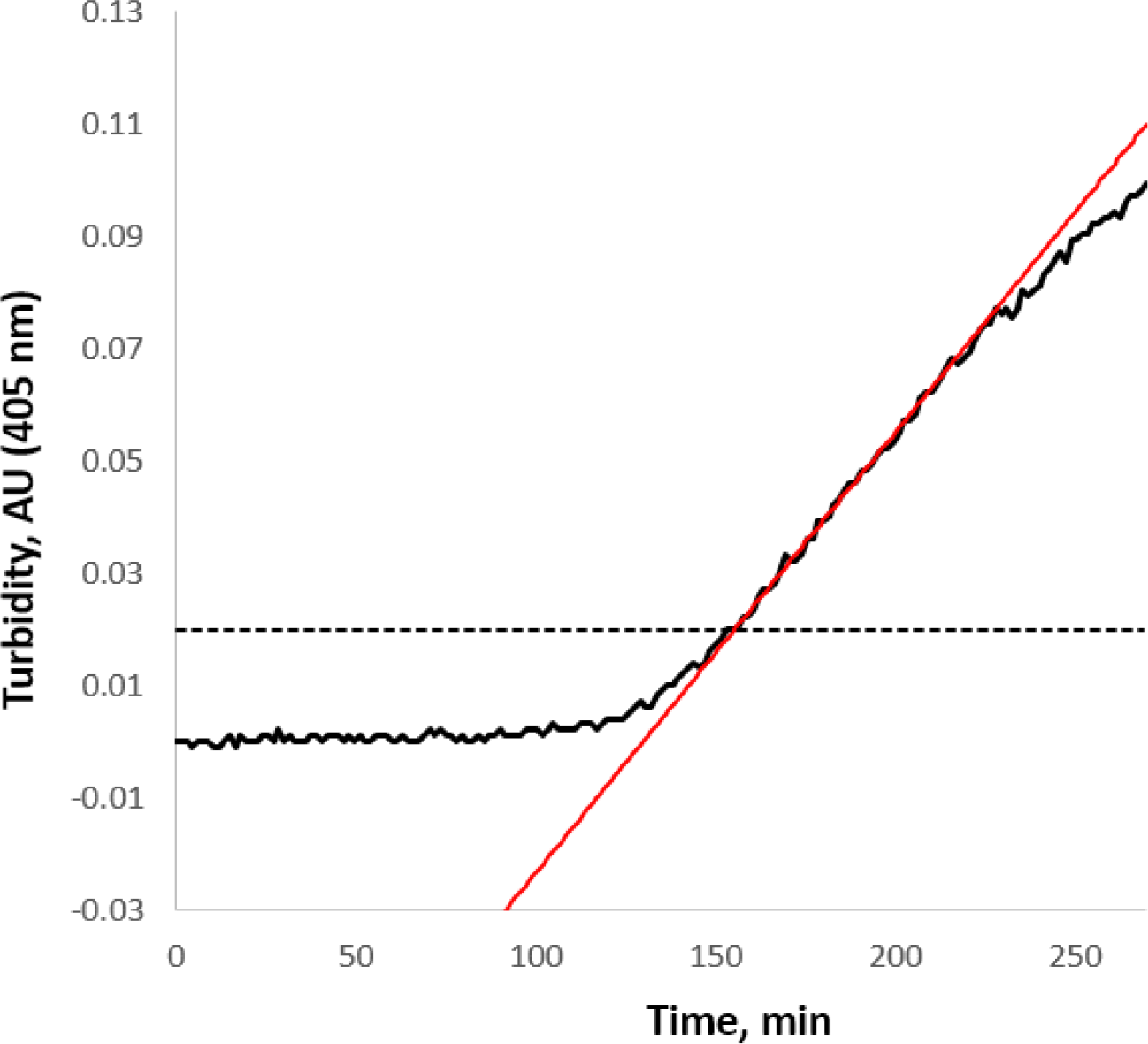
Example of maximum tangent and absolute turbidity threshold. A representative turbidity trace is shown for the aggregation at 37 °C and pH 7 of 20 μM W42Q with 70 μM WT serving as conformational catalyst. A substantial portion of the observed turbidity range features an approximately linear rate of increase. The maximum slope is determined by averaging slopes of linear regression fits of 20 sliding windows of 10 data points (15 min) each (45 min of data in total) immediately subsequent to the crossing of the turbidity threshold, set here at a constant value of 0.02 for the entire dataset.

**Fig. S6:**
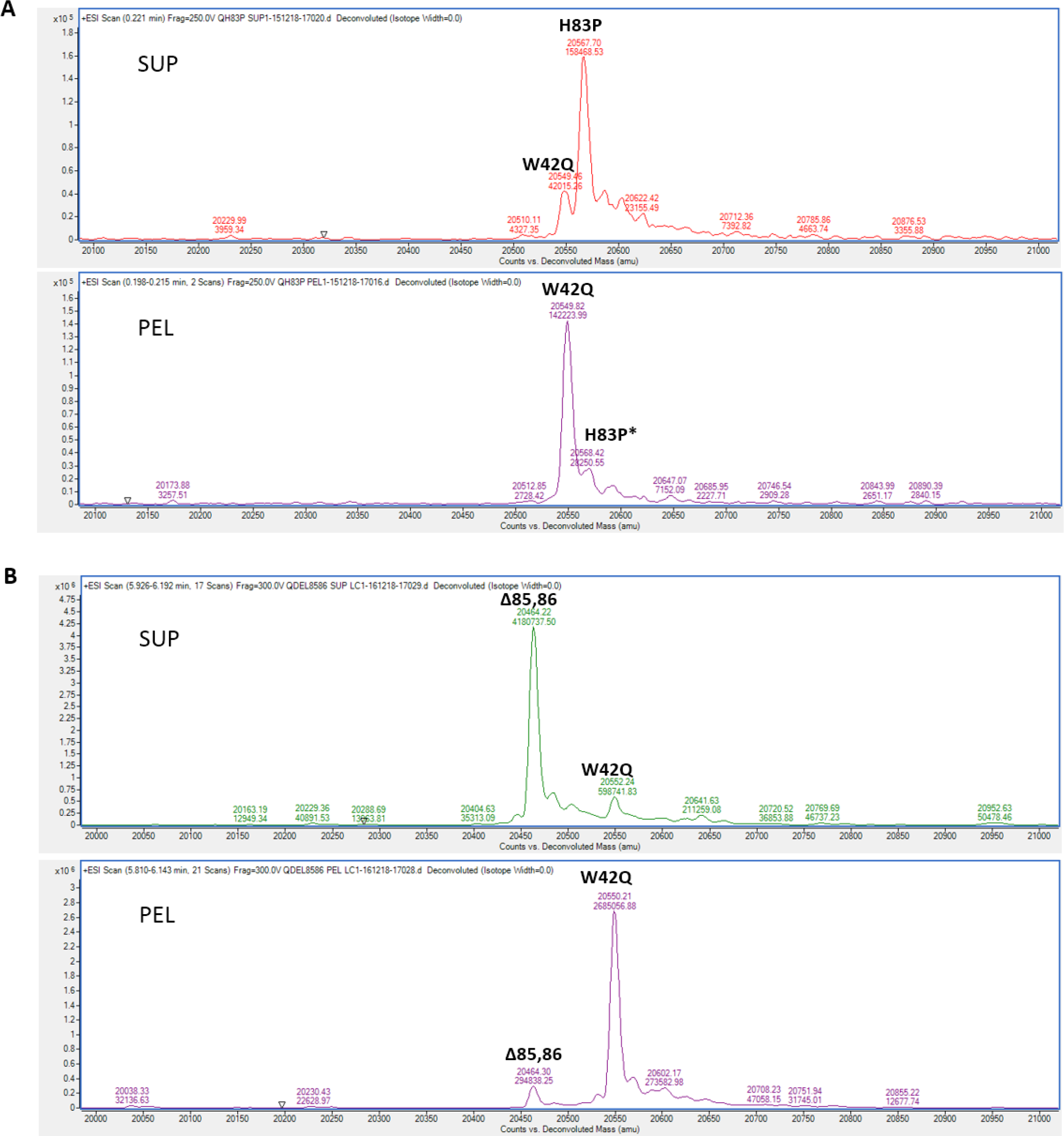
Coaggregation was observed between W42Q and ∆85,86. (*B*), but not between W42Q and H83P (*A*) in deconvoluted whole-protein electrospray mass spectra. Supernatant (“SUP”) and pellet (“PEL”) fractions from each aggregation experiment were analyzed. *Since their masses are too similar, the W42Q single oxidation (+16 Da.) peak and the native H83P peak cannot be reliably resolved on the Agilent 6220 instrument. However, comparison of the peak possibly containing H83P in (A) to the +16 peak in the PEL sample in (*B*), where no H83P is present, suggests that any co-aggregation of W42Q with H83P is at or below the limit of detection.

**Table S7:** Solvent-accessible hydrophobic surface area of HγD WT compared to its isolated domains and to BSA.

| Protein | SASA, Å<sup>2</sup> |
| --- | --- |
| HyD WT | 474 |
| HyD N-terminal domain | 1126 |
| HyD C-terminal domain | 971 |
| BSA | 5415 |

Visual Molecular Dynamics (VMD) software (U. Illinois) was used to calculate the sum of solvent-accessible surface area belonging to W, F, L, I, V, M, A, or P residues in HyD WT (PDB ID 1hk0), its isolated domains (by removing the cognate domain from the same PDB structure), and bovine serum albumin (BSA) monomer (PDB ID 3v03).

**Fig. S8:**
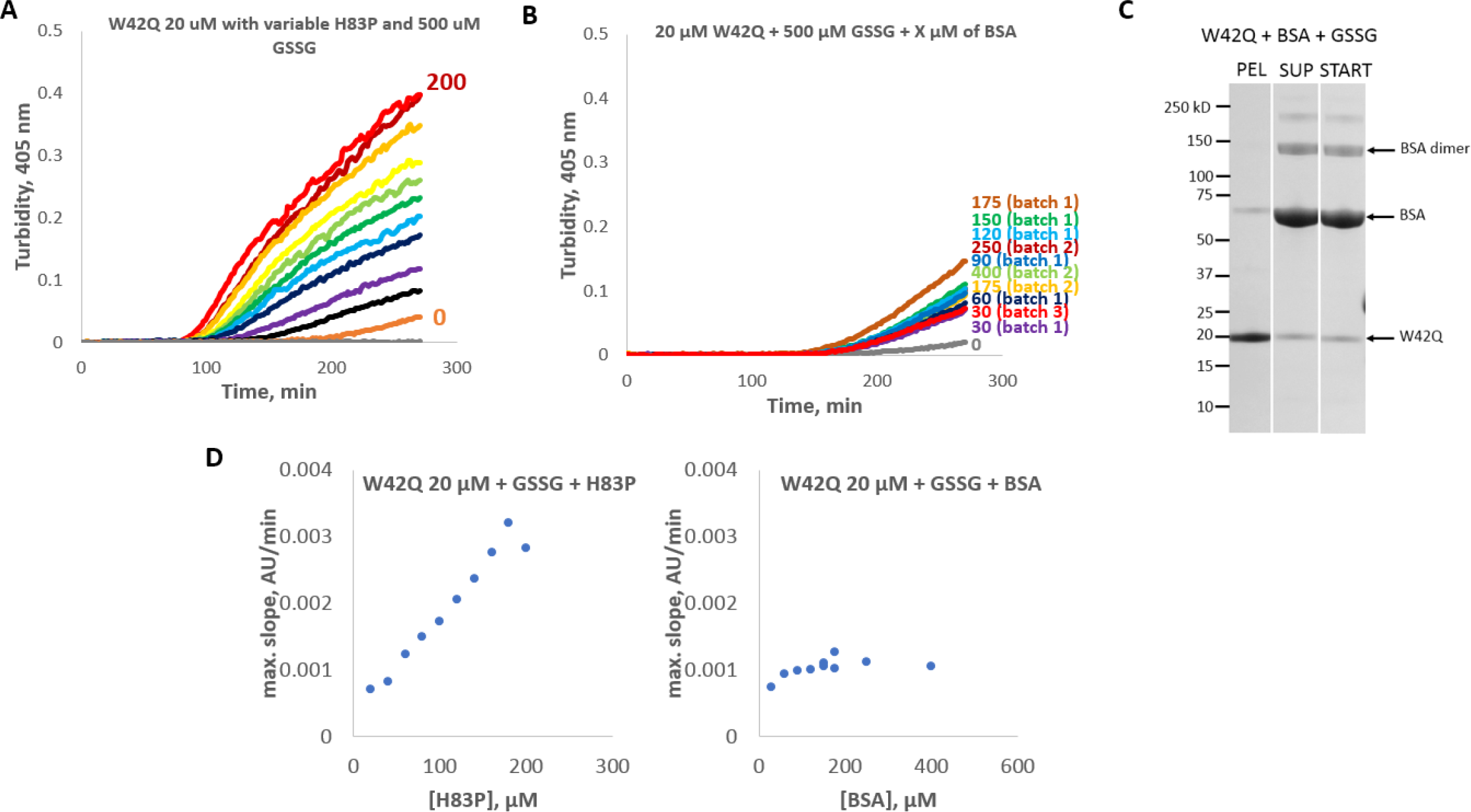
Conformational catalysis is not due to surface hydrophobicity. (*A*) Aggregation traces of W42Q with variable concentration (from 0 to 200 μM, with a step of 20 μM) of H83P revealed much greater turbidity than (*B*) the same in the presence of variable [BSA] from three separate batches as indicated. The BSA protein was prepared each time by dissolving several milligrams of BSA powder (Sigma), followed by size-exclusion chromatography, isolation and concentration of the monomer fraction. (*C*) Modest co-aggregation was observed between BSA and W42Q by pellet/supernatant separation. This may be due to disulfide-mediated cross-linking between the two proteins. (*D*) While presence of BSA did accelerate the maximum aggregation rate of W42Q, the effect was both smaller and much less dose-dependent than for H83P.

**Fig. S9:**
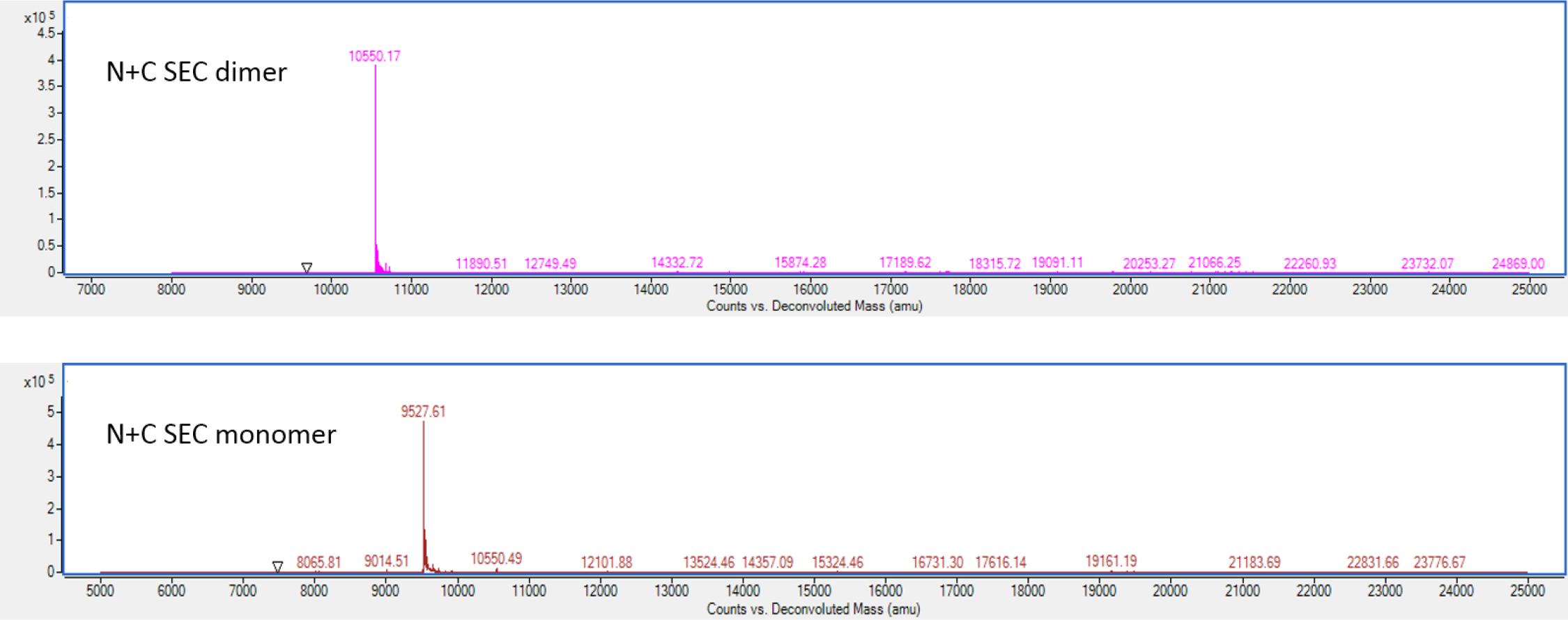
No heterodimers form in mixtures of isolated N-td and C-td. Deconvoluted HPLC/electrospray ionization mass spectrometry total ion intensities are shown for the two eluate peaks from size-exclusion chromatography of mixtures of isolated N-td and C-td in Fig. 4D. The first peak, a non-covalent dimer, returned only a peak that matched the expected molecular weight for the C-td (10,552 Da.). The second, monomeric peak returned only the expected molecular weight of the N-td (9528 Da.), with a trace amount of C-td likely arising from incomplete resolution of the two size-exclusion fractions. No covalently bonded dimers were detectable.

**Fig. S10:**
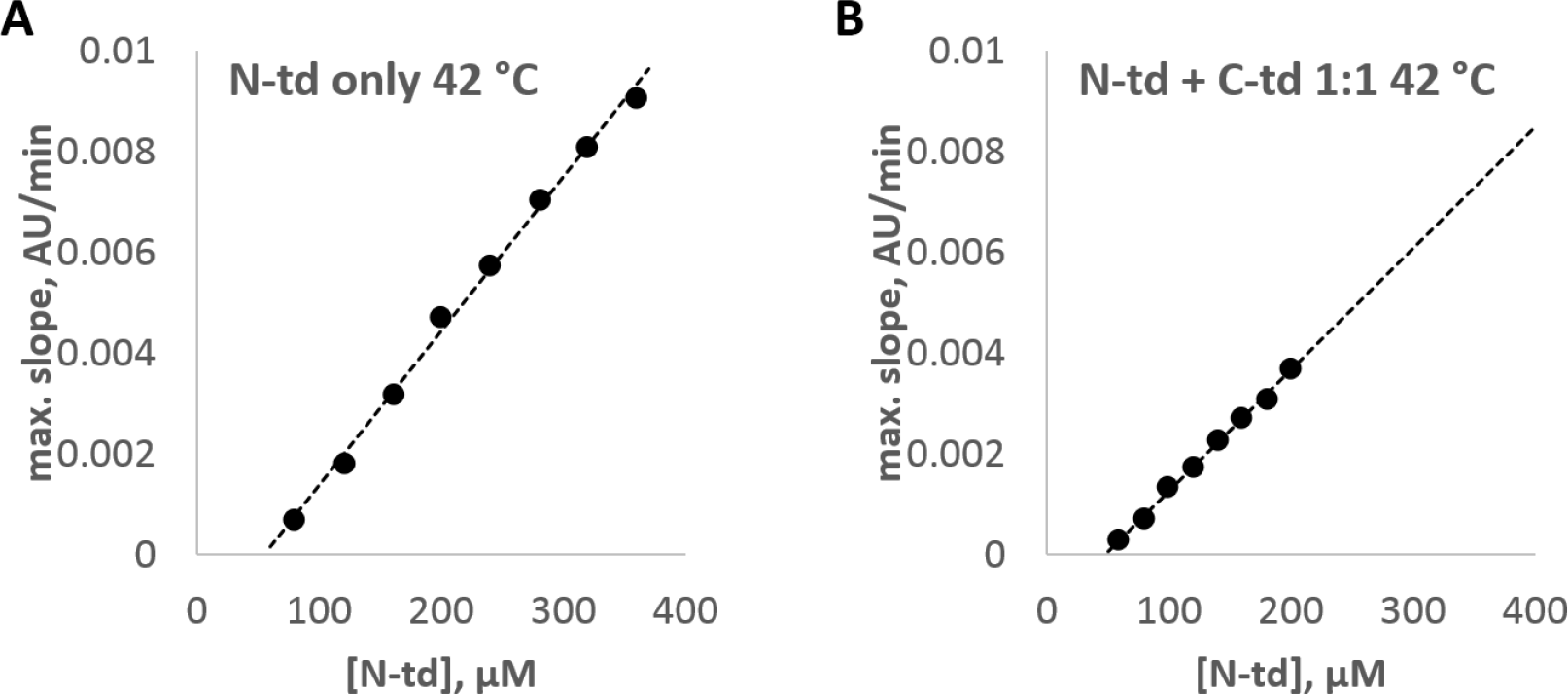
Isolated N-td shows a linear concentration dependence of maximal turbidity slope, regardless of presence of C-td. The isolated, unmutated N-terminal domain of HγD was incubated at 42 °C and pH 7 in the presence of 500 μM GSSG either by itself (A) or as an equimolar mixture with the isolated C-terminal domain (B). The elevated temperature was used since the concentration range of aggregation is impractically high at 37 °C for this construct. In each case, the maximal turbidity slope varied linearly with protein concentration, in sharp contrast to the quadratic concentration dependences observed for aggregation of the full-length L5S, W42Q, and V75D variants where the domains are physically linked to each other.

**Fig. S11:**
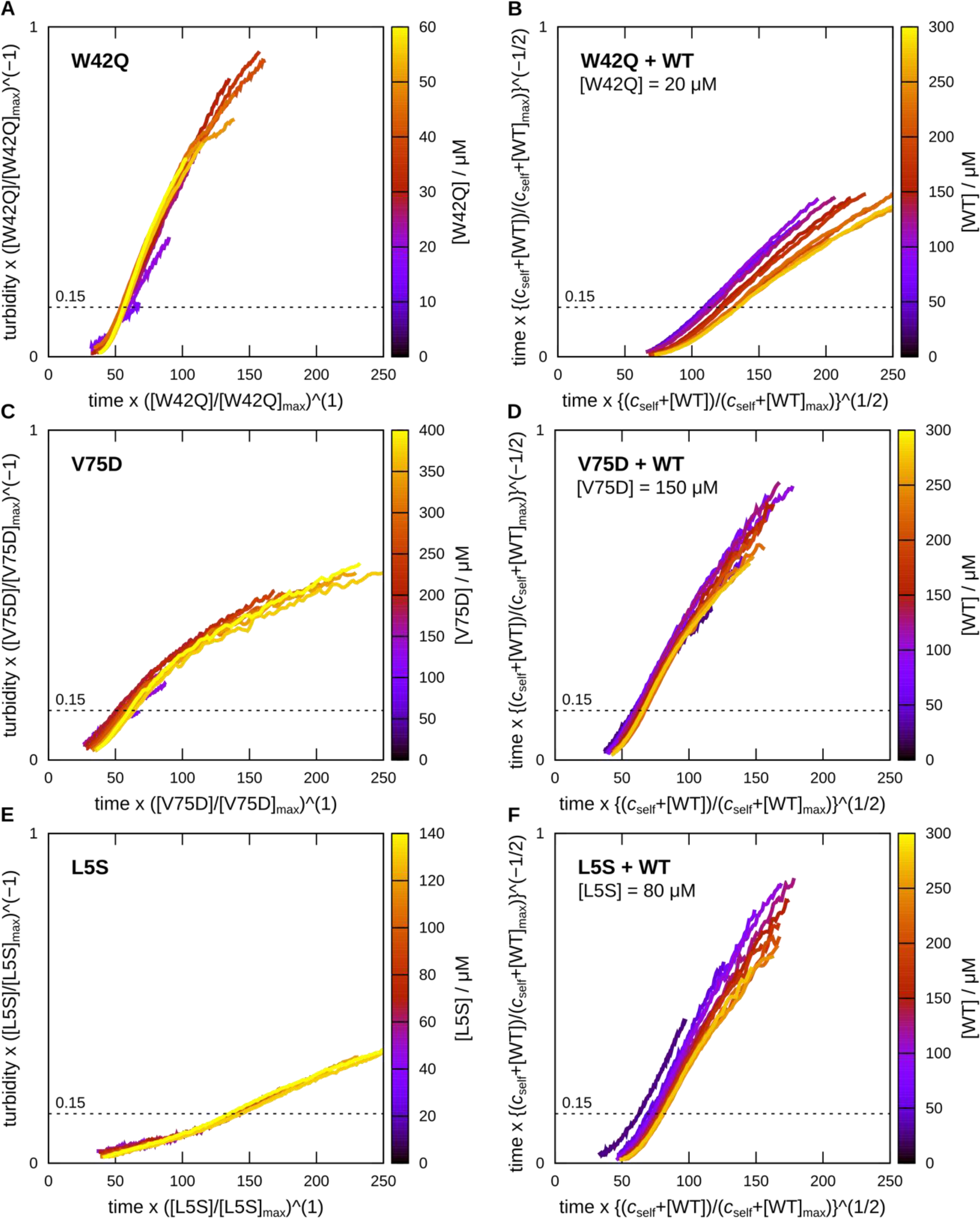
Turbidity traces scale with protein concentrations as predicted by the kinetic model. Time has units of minutes, and turbidity is reported in arbitrary units on a linear scale. The scaled turbidity threshold 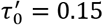 is indicated by a dotted line in each plot and used for all calculations shown in Fig. 6. (A,C,E) Self-catalysis: *J*_SS_~[N]^2^. (B,D,F) Heterogeneous catalysis: *J*_SS_~[N^WT^] + c_self_. The values for c_self_ (88 *μ*M, 33 *μ*M, and 13 *μ*M for the mutants W42Q, V75D, and L5S, respectively) were determined from a linear regression to the rate data shown in Fig. 6D using WT concentrations up to 150 *μ*M, where depletion effects tend to be smaller.

**Fig. S12:**
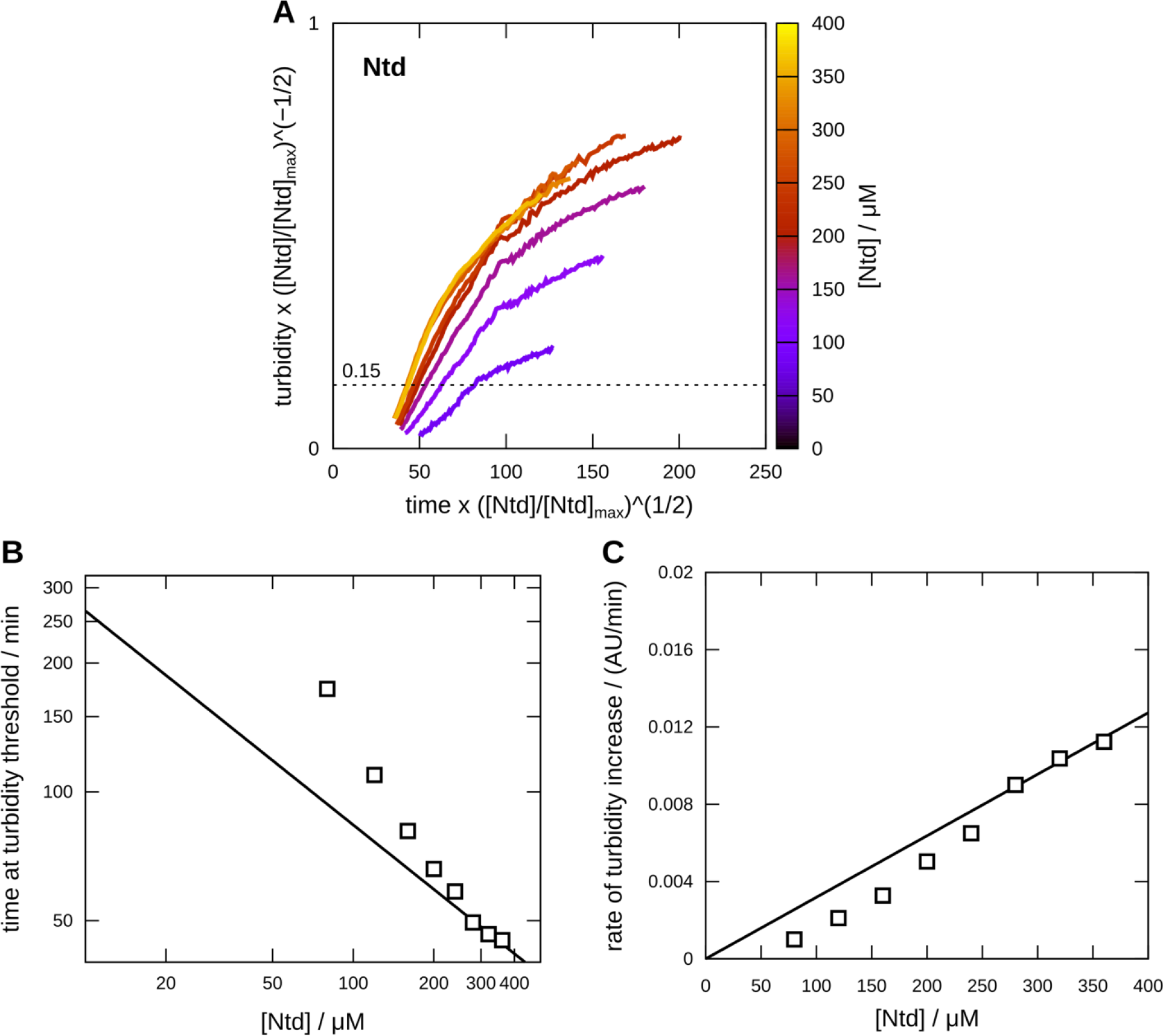
Analysis of turbidity data for the N-terminal domain with no catalyst. (A) Scaling analysis of non-catalyzed, unimolecular misfolding (cf. Fig. S11), with *J*_SS_~[N]. (B) The time until reaching a fixed scaled turbidity threshold is shown on a log—log plot. (C) The slope of the turbidity trace at a constant scaled time is shown on a linear plot. Solid lines show the ideal scaling behavior predicted by the aggregation scheme presented in SI Equation 1, fit to the highest concentration data. Within the framework of this model, deviations from the ideal scaling prediction may indicate that the misfolding time constant (inverse misfolding rate) is shorter than the duration of the experiment, leading to depletion effects that become more apparent as the concentration is lowered. Alternatively, the relatively poor fit to the ideal scaling prediction may indicate that a process that is not accounted for in SI Equation 1 plays a role in N-terminal domain aggregation, but not in the aggregation of the other mutants.

### SI Text

#### Kinetic model and scaling prediction

Assuming that the solution is well mixed, we consider an extremely general description of the aggregation of I_ss_ monomers [Smoluchowski 1916]:

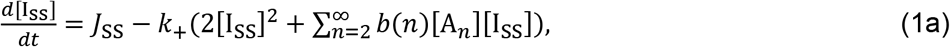

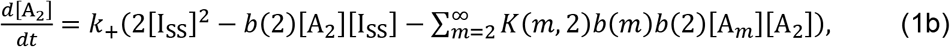

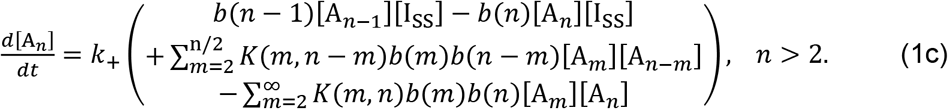

The minimum aggregate size is *n* = 2, *b*(*n*) is the number of binding sites on an aggregate of size *n* divided by the number of binding sites on a monomer, and *K* > 0 allows for coalescence of aggregates. Fragmentation and disassembly are not allowed because all aggregation events are assumed to be irreversible. At time *t* = 0, [I_ss_] and all aggregate concentrations are zero.

We initially assume that *J*_SS_ is a constant. Then, changing variables such that 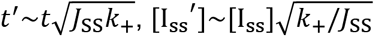, and 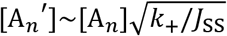, we find that the first equation becomes

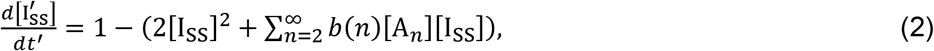

which is independent of *J*_SS_ and *k*_+_. Transformation of the equations for [A_*n*_] similarly eliminates the dependence on the rate constant *k*_+_. Consequently, if *J*_SS_ is a constant, then the distribution of aggregate concentrations at any time *t* is equal to the unique distribution of aggregation concentrations in the scaled model at time *t*′, multiplied by the concentration scaling ratio [I_ss_]/[I_ss_′]. We refer to this as the “ideal” scaling behavior.

#### Depletion effects

##### Unimolecular misfolding and heterogeneous catalysis

We first consider the case in which misfolding involves only one molecule of the aggregation-prone protein. In the case of unimolecular misfolding, the flux of misfolded protein is

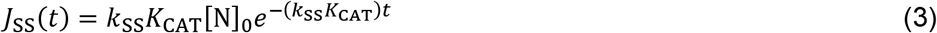

and we denote the initial concentration of the native-state mutant protein by [N]_0_. Switching to the scaled variables, where 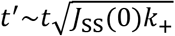 and so forth, Equation 2 becomes

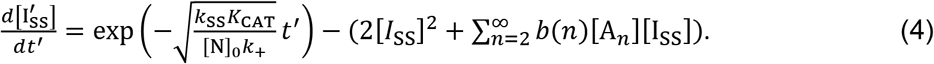

Thus, even in the scaled variables, the aggregation process is dependent on the initial concentration of native-state protein. The turbidity traces should only converge to a master curve in the limit of a high initial concentration and a slow misfolding rate.

In the case of heterogeneous catalysis, we model the contribution of WT-catalyzed misfolding by replacing *K*_CAT_ in Equation 4 with 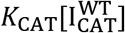. Deviations from a master turbidity curve are therefore dependent on the concentration of the WT protein, with the effect being greater at high WT concentrations and low aggregation-prone protein concentrations.

##### Self-catalysis

In the case of bi-molecular self-catalysis, depletion leads to hyperbolic decay of the native protein concentration, so that the flux of misfolded protein is

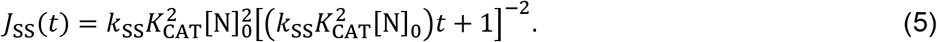

In terms of the scaled variables, Equation 2 becomes

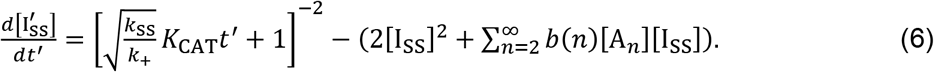

The effect of depletion in this case is *independent* of the initial native-state protein concentration [N]_0_. Therefore, even though depletion still affects the shape of the master turbidity curve, varying the protein concentration alone should not result in deviations from this curve after the scaling transformation has been applied.

## References

1. Han JH, Batey S, Nickson AA, Teichmann SA, & Clarke J (2007) The folding and evolution of multidomain proteins. Nat. Rev. Mol. Cell Biol. 8(4):319–330.

2. Kumar V & Chaudhuri TK (2018) Spontaneous refolding of the large multidomain protein malate synthase G proceeds through misfolding traps. J. Biol. Chem. 293(34):13270–13283.

3. Kantaev R, et al. (2018) Manipulating the Folding Landscape of a Multidomain Protein. J. Phys. Chem. B 122(49):11030–11038.

4. Borgia MB, et al. (2011) Single-molecule fluorescence reveals sequence-specific misfolding in multidomain proteins. Nature 474(7353):662–U142.

5. Borgia A, et al. (2015) Transient misfolding dominates multidomain protein folding. Nat. Commun. 6:10.

6. Brockwell DJ & Radford SE (2007) Intermediates: ubiquitous species on folding energy landscapes? Curr. Op. Struct. Biol. 17(1):30–37.

7. Bershtein S, Mu WM, Serohijos AWR, Zhou JW, & Shakhnovich EI (2013) Protein Quality Control Acts on Folding Intermediates to Shape the Effects of Mutations on Organismal Fitness. Mol. Cell 49(1):133–144.

8. Mitraki A, Fane B, Haase-Pettingell C, Sturtevant J, & King J (1991) Global suppression of protein folding defects and inclusion body formation. Science 253(5015):54–58.

9. Jacobs WM & Shakhnovich EI (2017) Evidence of evolutionary selection for cotranslational folding. Proc. Natl. Acad. Sci. U. S. A. 114(43):11434–11439.

10. Drummond DA & Wilke CO (2008) Mistranslation-induced protein misfolding as a dominant constraint on coding-sequence evolution. Cell 134(2):341–352.

11. Horwich A (2002) Protein aggregation in disease: a role for folding intermediates forming specific multimeric interactions. Journal of Clinical Investigation 110(9):1221–1232.

12. Chiti F & Dobson CM (2006) Protein misfolding, functional amyloid, and human disease. Annual Review of Biochemistry, Annual Review of Biochemistry), Vol 75, pp 333–366.

13. Chiti F & Dobson CM (2009) Amyloid formation by globular proteins under native conditions. Nat Chem Biol 5(1):15–22.

14. Wang Y, Papaleo E, & Lindorff-Larsen K (2016) Mapping transiently formed and sparsely populated conformations on a complex energy landscape. Elife 5:35.

15. Karamanos TK, Kalverda AP, Thompson GS, & Radford SE (2014) Visualization of transient protein-protein interactions that promote or inhibit amyloid assembly. Mol Cell 55(2):214–226.

16. Sun X, Dyson HJ, & Wright PE (2018) Kinetic analysis of the multistep aggregation pathway of human transthyretin. Proc. Natl. Acad. Sci. U. S. A. 115(27):E6201–E6208.

17. Serebryany E & King JA (2014) The βγ-crystallins: Native state stability and pathways to aggregation. Progress in Biophysics & Molecular Biology 115:32–41.

18. Bah A & Forman-Kay JD (2016) Modulation of Intrinsically Disordered Protein Function by Post-translational Modifications. J Biol Chem 291(13):6696–6705.

19. Sevcsik E, Trexler AJ, Dunn JM, & Rhoades E (2011) Allostery in a Disordered Protein: Oxidative Modifications to alpha-Synuclein Act Distally To Regulate Membrane Binding. J. Am. Chem. Soc. 133(18):7152–7158.

20. Schmitt ND & Agar JN (2017) Parsing disease-relevant protein modifications from epiphenomena: perspective on the structural basis of SOD1-mediated ALS. Journal of Mass Spectrometry 52(7):480–491.

21. Darling AL & Uversky VN (2018) Intrinsic Disorder and Posttranslational Modifications: The Darker Side of the Biological Dark Matter. Front. Genet. 9:18.

22. Deis LN, et al. (2014) Multiscale conformational heterogeneity in staphylococcal protein a: possible determinant of functional plasticity. Structure 22(10):1467–1477.

23. de Graff AM, Hazoglou MJ, & Dill KA (2016) Highly Charged Proteins: The Achilles’ Heel of Aging Proteomes. Structure 24(2):329–336.

24. Lampi KJ, Wilmarth PA, Murray MR, & David LL (2014) Lens beta-crystallins: the role of deamidation and related modifications in aging and cataract. Prog Biophys Mol Biol 115(1):21–31.

25. Bloemendal H, et al. (2004) Ageing and vision: structure, stability and function of lens crystallins. Prog Biophys Mol Biol 86(3):407–485.

26. Wride MA (2011) Lens fibre cell differentiation and organelle loss: many paths lead to clarity. Philos Trans R Soc Lond B Biol Sci 366(1568):1219–1233.

27. Nielsen J, et al. (2016) Eye lens radiocarbon reveals centuries of longevity in the Greenland shark (Somniosus microcephalus). Science 353(6300):702–704.

28. Kosinski-Collins MS & King J (2003) In vitro unfolding, refolding, and polymerization of human gamma D crystallin, a protein involved in cataract formation. Protein Sci. 12(3):480–490.

29. Moreau KL & King JA (2012) Protein misfolding and aggregation in cataract disease and prospects for prevention. Trends Mol. Med. 18(5):273–282.

30. Truscott RJW, Schey KL, & Friedrich MG (2016) Old Proteins in Man: A Field in its Infancy. Trends Biochem Sci 41(8):654–664.

31. Ma Z, et al. (1998) Age-related changes in human lens crystallins identified by HPLC and mass spectrometry. Exp Eye Res 67(1):21–30.

32. Friedburg D & Manthey KF (1973) Glutathione and NADP linked enzymes in human senile cataract. Exp Eye Res 15(2):173–177.

33. Sweeney MH & Truscott RJ (1998) An impediment to glutathione diffusion in older normal human lenses: a possible precondition for nuclear cataract. Exp Eye Res 67(5):587–595.

34. Hains PG & Truscott RJ (2008) Proteomic analysis of the oxidation of cysteine residues in human age-related nuclear cataract lenses. Biochim Biophys Acta 1784(12):1959–1964.

35. Fan X, et al. (2015) Evidence of Highly Conserved β-Crystallin Disulfidome that Can be Mimicked by In Vitro Oxidation in Age-related Human Cataract and Glutathione Depleted Mouse Lens. Mol. Cell. Proteomics 14(12):3211–3223.

36. Wang B, et al. (2017) The oxidized thiol proteome in aging and cataractous mouse and human lens revealed by ICAT labeling. Aging Cell 16(2):244–261.

37. Ramkumar S, Fan X, Wang B, Yang S, & Monnier VM (2018) Reactive cysteine residues in the oxidative dimerization and Cu2+ induced aggregation of human γD-crystallin: Implications for age-related cataract. Biochim Biophys Acta 1864:3595–3604.

38. Lou MF (2003) Redox regulation in the lens. Progress in Retinal and Eye Research 22(5):657–682.

39. Takemoto LJ (1996) Oxidation of cysteine residues from alpha-A crystallin during cataractogenesis of the human lens. Biochem Biophys Res Commun 223(2):216–220.

40. Takemoto LJ (1997) Disulfide bond formation of cysteine-37 and cysteine-66 of beta B2 crystallin during cataractogenesis of the human lens. Exp Eye Res 64(4):609–614.

41. Takemoto LJ (1997) BetaA3/A1 crystallin from human cataractous lens contains an intramolecular disulfide bond. Current Eye Research 16(7):719–724.

42. Hanson SR, Smith DL, & Smith JB (1998) Deamidation and disulfide bonding in human lens gamma-crystallins. Exp Eye Res 67(3):301–312.

43. Serebryany E, et al. (2016) An Internal Disulfide Locks a Misfolded Aggregation-prone Intermediate in Cataract-linked Mutants of Human γD-Crystallin. J. Biol. Chem. 291(36):19172–19183.

44. Serebryany E, Yu S, Trauger SA, Budnik B, & Shakhnovich EI (2018) Dynamic disulfide exchange in a crystallin protein in the human eye lens promotes cataract-associated aggregation. J Biol Chem In press.

45. Serebryany E, et al. (2016) Aggregation of Trp > Glu point mutants of human gamma-D crystallin provides a model for hereditary or UV-induced cataract. Protein Sci 25(6):1115–1128.

46. Tielsch JM, Kempen JH, Congdon N, & Friedman DS (2008) Vision probelms in the U.S.: Prevalence of adult visual impairment and age-related eye disease in America. Prevent Blindness America.

47. Liu YC, Wilkins M, Kim T, Malyugin B, & Mehta JS (2017) Cataracts. Lancet 390(10094):600–612.

48. Graw J (2009) Genetics of crystallins: Cataract and beyond. Exp. Eye Res. 88(2):173–189.

49. Vendra VPR, Khan I, Chandani S, Muniyandi A, & Balasubramanian D (2016) Gamma crystallins of the human eye lens. Biochim. Biophys. Acta-Gen. Subj. 1860(1):333–343.

50. Shiels A & Hejtmancik JF (2017) Mutations and mechanisms in congenital and age-related cataracts. Experimental Eye Research 156:95–102.

51. Chew FLM, et al. (2018) Estimates of visual impairment and its causes from the National Eye Survey in Malaysia (NESII). PLoS One 13(6).

52. Basak A, et al. (2003) High-resolution X-ray crystal structures of human gamma D crystallin (1.25 angstrom) and the R58H mutant (1.15 angstrom) associated with aculeiform cataract. J. Mol. Biol. 328(5):1137–1147.

53. Flaugh SL, Kosinski-Collins MS, & King J (2005) Contributions of hydrophobic domain interface interactions to the folding and stability of human gammaD-crystallin. Protein Sci 14(3):569–581.

54. Flaugh SL, Mills-Henry IA, & King J (2006) Glutamine deamidation destabilizes human gammaD-crystallin and lowers the kinetic barrier to unfolding. J. Biol. Chem. 281(41):30782–30793.

55. Mills-Henry IA, Flaugh SL, Kosinski-Collins MS, & King JA (2007) Folding and stability of the isolated Greek key domains of the long-lived human lens proteins gammaD-crystallin and gammaS-crystallin. Protein Sci 16(11):2427–2444.

56. Serebryany E (2016) Mechanism of aggregation of oxidation-mimicking mutants of human gamma-D crystallin. Ph.D. (Massachusetts Institute of Technology).

57. Serebryany E & King JA (2015) Wild-type human gammaD-crystallin promotes aggregation of its oxidation-mimicking, misfolding-prone W42Q mutant. J Biol Chem 290(18):11491–11503.

58. Ji FL, Jung J, Koharudin LMI, & Gronenborn AM (2013) The Human W42R gamma D-Crystallin Mutant Structure Provides a Link between Congenital and Age-related Cataracts. J. Biol. Chem. 288(1):99–109.

59. Sinha D, et al. (2001) A temperature-sensitive mutation of Crygs in the murine Opj cataract. J. Biol. Chem. 276(12):9308–9315.

60. Graw J, et al. (2002) V76D mutation in a conserved gamma D-crystallin region leads to dominant cataracts in mice. Mammalian Genome 13(8):452–455.

61. Wang BB, et al. (2011) A Novel CRYGD Mutation (p.Trp43Arg) Causing Autosomal Dominant Congenital Cataract in a Chinese Family. Human Mutation 32(1):E1939–E1947.

62. Wong EK (2018) Modeling the structure and dynamics of gamma-crystallins and their cataract-related variants. Ph.D. (University of California, Irvine).

63. Woodard JC (2017) Monte Carlo simulation approaches to protein stability and aggregation prediction. Ph.D. (Harvard University, Cambridge, MA, USA).

64. Jung J, Byeon IJL, Wang YT, King J, & Gronenborn AM (2009) The Structure of the Cataract-Causing P23T Mutant of Human gamma D-Crystallin Exhibits Distinctive Local Conformational and Dynamic Changes. Biochemistry 48(12):2597–2609.

65. Mills-Henry IA, Thol SL, Kosinski-Collins MS, Serebryany E, & King JA (2019) Kinetic stability of long-lived human gamma-D and gamma-S crystallins and their isolated double Greek Key domains. Biophys. J. In Revision.

66. Purkiss AG, Bateman OA, Goodfellow JM, Lubsen NH, & Slingsby C (2002) The x-ray crystal structure of human gamma S-crystallin C-terminal domain. J. Biol. Chem. 277(6):4199–4205.

67. Flyvbjerg H, Jobs E, & Leibler S (1996) Kinetics of self-assembling microtubules: An “inverse problem” in biochemistry. Proc. Natl. Acad. Sci. U. S. A. 93(12):5975–5979.

68. Ray NJ, Hall D, & Carver JA (2016) Deamidation of N76 in human gamma S-crystallin promotes dimer formation. Biochim. Biophys. Acta 1860(1):315–324.

69. Lubsen NH, Aarts HJM, & Schoenmakers JGG (1988) The Evolution of Lenticular Proteins -- The Beta-crystallin and Gamma-crystallin Super Gene Family. Prog. Biophys. Mol. Biol. 51(1):47–76.

70. Lukatsky DB, Shakhnovich BE, Mintseris J, & Shakhnovich EI (2007) Structural similarity enhances interaction propensity of proteins. J. Mol. Biol. 365(5):1596–1606.

71. Lukatsky DB, Zeldovich KB, & Shakhnovich EI (2006) Statistically enhanced self-attraction of random patterns. Phys. Rev. Lett. 97(17).

72. Wright CF, Teichmann SA, Clarke J, & Dobson CM (2005) The importance of sequence diversity in the aggregation and evolution of proteins. Nature 438(7069):878–881.

73. Thorn DC, et al. (2019) The Structure and Stability of the Disulfide-Linked gamma S-Crystallin Dimer Provide Insight into Oxidation Products Associated with Lens Cataract Formation. J. Mol. Biol. 431(3):483–497.

74. Srivastava OP & Srivastava K (1998) Degradation of gamma D- and gamma s-crystallins in human lenses. Biochem. Biophys. Res. Commun. 253(2):288–294.

75. Manhart M & Morozov AV (2015) Protein folding and binding can emerge as evolutionary spandrels through structural coupling. Proc. Natl. Acad. Sci. U. S. A. 112(6):1797–1802.

76. Dixit PD & Maslov S (2013) Evolutionary Capacitance and Control of Protein Stability in Protein-Protein Interaction Networks. PLoS Comp. Biol. 9(4).

77. Aguzzi A & Calella AM (2009) Prions: protein aggregation and infectious diseases. Physiol Rev 89(4):1105–1152.

78. Tessier PM & Lindquist S (2007) Prion recognition elements govern nucleation, strain specificity and species barriers. Nature 447(7144):556–561.

79. Grad LI & Cashman NR (2014) Prion-like activity of Cu/Zn superoxide dismutase: implications for amyotrophic lateral sclerosis. Prion 8(1):33–41.

80. Slingsby C & Wistow GJ (2014) Functions of crystallins in and out of lens: Roles in elongated and post-mitotic cells. Prog. Biophys. Mol. Biol. 115(1):52–67.

81. Quintanar L, et al. (2016) Copper and Zinc Ions Specifically Promote Nonamyloid Aggregation of the Highly Stable Human gamma-D Crystallin. ACS Chem Biol 11(1):263–272.

82. Dominguez-Calva JA, Perez-Vazquez ML, Serebryany E, King JA, & Quintanar L (2018) Mercury-induced aggregation of human lens gamma-crystallins reveals a potential role in cataract disease. J. Biol. Inorg. Chem. 23(7):1105–1118.

83. Dominguez-Calva JA, Haase-Pettingell C, Serebryany E, King JA, & Quintanar L (2018) A Histidine Switch for Zn-Induced Aggregation of gamma-Crystallins Reveals a Metal-Bridging Mechanism That Is Relevant to Cataract Disease. Biochemistry 57(33):4959–4962.

84. Keller BO & Li L (2006) Three-layer matrix/sample preparation method for MALDI MS analysis of low nanomolar protein samples. J. Am. Soc. Mass Spectr. 17(6):780–785.

## SI Reference

Von Smoluchowski, Marian. “Drei vortrage uber diffusion. Brownsche bewegung und koagulation von kolloidteilchen.” Z. Phys. 17 (1916): 557–585.

